# Contractile fibroblasts are recruited to the growing mammary epithelium to support branching morphogenesis

**DOI:** 10.1101/2024.06.05.597593

**Authors:** Jakub Sumbal, Robin P. Journot, Marisa M. Faraldo, Zuzana Sumbalova Koledova, Silvia Fre

## Abstract

Fibroblasts are stromal cells found in connective tissue that are critical for organ development, homeostasis, and disease. Single-cell transcriptomic analyses have revealed a high level of inter- and intra-organ heterogeneity of fibroblasts. However, the functional implications and lineage relations of different fibroblast subtypes remain unexplored, especially in the mammary gland. Here we provide a comprehensive characterization of pubertal mammary fibroblasts, achieved using single-cell RNA sequencing, spatial mapping, and in vivo lineage tracing. Notably, we discovered a transient niche-forming population of specialized contractile fibroblasts that exclusively localize around the tips of the growing mammary epithelium and are recruited from the surrounding fat pad. Using functional organoid-fibroblast co-cultures we reveal that different fibroblast populations can acquire contractile features when in direct contact with the epithelium, promoting morphogenesis. In summary, our exhaustive characterization of these specialized cells provides new insights into mammary fibroblast heterogeneity and implicates their functional relevance for branching morphogenesis and lineage hierarchy during mouse mammary gland development.

## Introduction

Fibroblasts are cells of mesenchymal origin that form the soft connective tissue, and they are extremely pleiotropic in their function. They secrete, remodel, and degrade components of the extracellular matrix (ECM), produce signaling molecules to communicate with other stromal cells, immune cells or organ parenchyma, act as a source of other mesenchymal cells and can become contractile and generate mechanical forces within tissues^1–3^. Through a delicate spatiotemporal orchestration of the aforementioned functions, fibroblasts guide development, maintain homeostasis and support pathological changes in many organs^1^. Recent studies employing single-cell transcriptomics suggest that fibroblasts represent a heterogenous cell population with distinct cell types or cell states specialized for different functions, presenting inter-organ conserved as well as divergent behavior and functions^4,5^. It is not yet understood, however, how fibroblast heterogeneity is linked to their plasticity and their origin.

Epithelial ductal elongation and branching in the mammary gland occurs mostly postnatally, when a peak of estrogen during puberty awakens the rudimentary mammary epithelium formed during embryogenesis, and the mammary epithelium actively proliferates and invades the surrounding fat pad stroma to generate a highly branched ductal network^6^. This morphogenetic function is executed by terminal end buds (TEBs), bulb-shaped and highly proliferative epithelial structures that form at the tips of the ducts and invade the surrounding stroma at the astonishing speed of 0.5 mm per day^7^. In a stochastic pattern, TEBs bifurcate to generate new branches^8,9^. The cellular and physical mechanisms that drive *in vivo* branching morphogenesis, TEB bifurcation and ductal elongation have not been thoroughly explored.

The importance of the stroma, in particular of fibroblasts, to support mammary branching morphogenesis is well recognized^2,10^. Mammary fibroblasts secrete growth factors^11^ that can induce epithelial branching^12^. Fibroblasts also secrete and remodel collagen 1^13^ that supports ductal elongation in organoids^14^. Likewise, *in vivo* collagen deposition correlates with the direction of ductal elongation^15^ and with TEB bifurcations^16^. Importantly, we have recently demonstrated that contractile mammary fibroblasts are instrumental to the induction of mammary epithelial organoid branching^17^, similarly to smooth muscle cells in the budding embryonic lung^18^. We have also reported the presence of contractile α-smooth muscle actin (αSMA, encoded by *Acta2*) positive (αSMA+) fibroblasts *in vivo,* around TEBs in the pubertal mammary gland undergoing branching morphogenesis^17^. Fibroblast contractility is an important marker of their biology, often linked with their function and activation state during organ development^18,19^, wound healing, fibrosis^20^ as well as in cancer^21,22^.

The transcriptional and functional heterogeneity of mammary fibroblasts during the prominent epithelial remodeling underlying pubertal branching morphogenesis *in vivo* has not been explored. Although single cell RNA sequencing (scRNAseq) datasets of mammary fibroblasts were generated^23–26^, these studies only analyzed the mammary gland under homeostatic conditions and lacked spatial analysis. In this study, we have rigorously characterized the diversity of pubertal fibroblasts and identified a specialized subset of spatially restricted contractile fibroblasts that exist only in proximity of growing TEBs. We have thus analyzed their behavior, function and relation to other mammary fibroblasts using scRNAseq, spatial mapping, transplantations, in vitro co-cultures with organoids and in vivo lineage tracing approaches.

## Results

### Single cell analysis of the growing tips and the subtending ductal regions of the pubertal mammary gland reveals the existence of contractile peri-TEB fibroblasts

TEBs of developing mammary glands are surrounded by a stromal niche with a thickened peri-epithelial fibroblast layer^27^ (Figure 1A, B), and we recently found that peri-TEB fibroblasts express the contractility marker αSMA (Figure 1C, Movie 1)^17^. To characterize the αSMA+ peri-TEB fibroblasts in more depth, we performed scRNAseq on cells dissociated from micro-dissected regions of pubertal mammary glands, containing either mainly TEBs or subtending ducts (Figure 1D). We purposedly chose not to FACS sort cells to prevent sample biasing and proceeded to a comparative analysis of the cell types derived from the two distinct anatomical sites, one enriched in TEBs (hence in peri-TEB fibroblasts) and one deprived of them. In both samples, we identified distinct fibroblast clusters based on generic markers (*Col1a1, Col1a2, Pdgfra,* Supplementary figure 1A, B) and merged them to create a collective UMAP graph representing all pubertal mammary fibroblasts (Figure 1E, F). Of note, our dataset did not contain pericytes (Supplementary figure 1B) but included epithelial cells (Supplementary figure 1C-E) and leukocytes, mainly T cells, B cells, macrophages, and dendritic cells (Supplementary figure 1F-H).

**Fig. 1.**
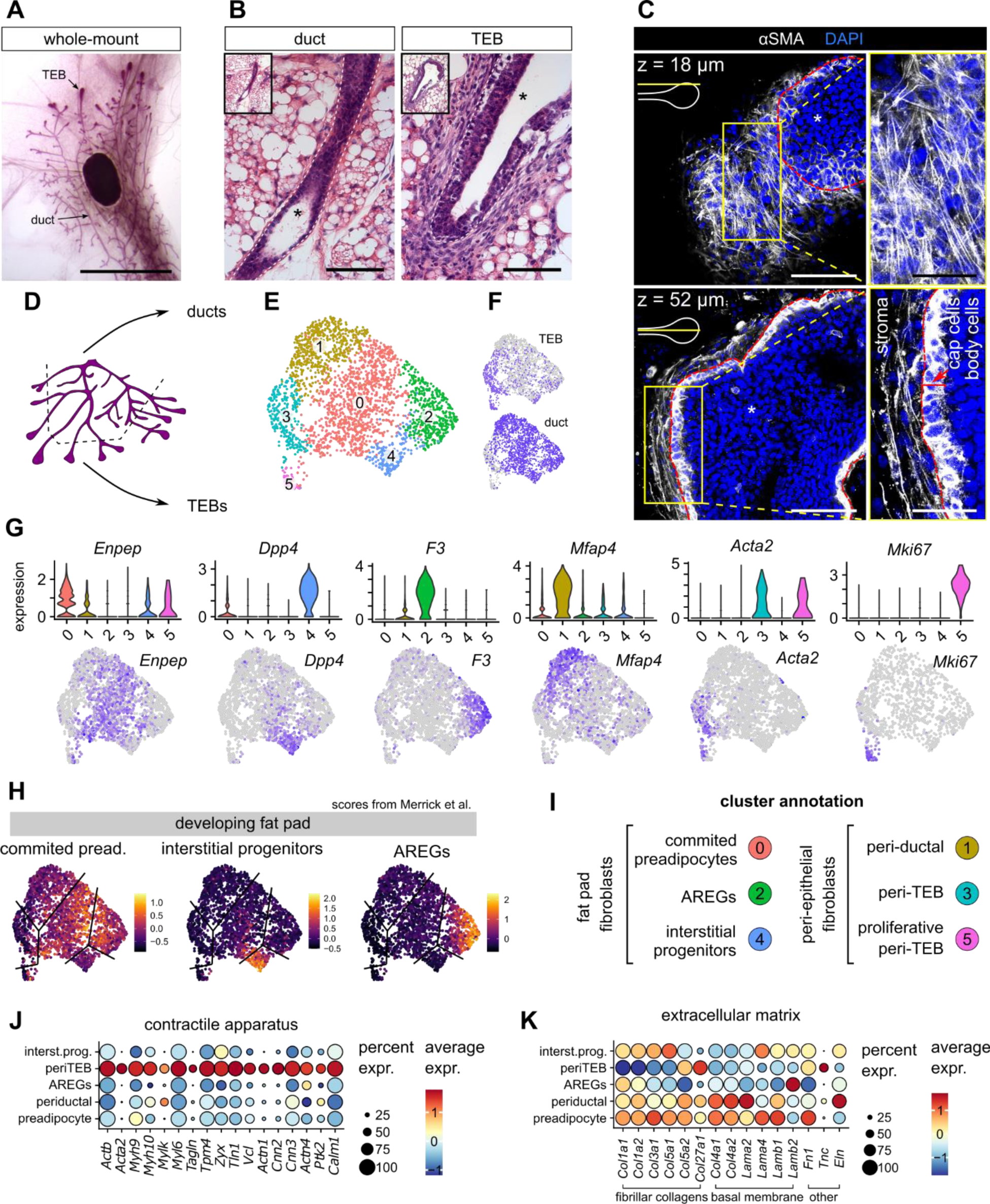
Single cell RNA sequencing of pubertal mammary fibroblasts. **A.** A representative image of a pubertal mammary gland from a 5 week old mouse. Scale bar: 1 cm. **B.** Histological detail of epithelial duct and TEB. Scale bar: 100 µm. Ep. marks epithelial compartment. **C.** Immunostaining for αSMA in a cleared mammary gland detects fibroblasts around TEB (TEB borders are delineated by a red line) as well as cap cells of the TEB. Top and middle optical sections of the imaged z stack are presented; their localization according to TEB is indicated by yellow line. The full z-stack is presented in Movie 1. Scale bars: 100 µm, 50 µm in detail. Ep. marks epithelium. **D.** A scheme of microdissected regions of the mammary gland. **E-F.** UMAP of merged mammary fibroblasts showing fibroblast clusters (**E**) and sample of origin (**F**). **G.** Violin plots and UMAPs showing expression of cluster marker genes. **H.** Transcriptomic scores of fibroblasts from developing fat pad (Merrick et al. 2019) plotted over our dataset of mammary fibroblasts. The black lines indicate approximate borders between our 6 clusters. **I.** Annotation of fibroblast clusters. **J, K.** Dot plots showing expression of contractility-related genes (**J**) and genes encoding extracellular matrix (**K**).

Unsupervised clustering of merged fibroblasts identified six separate clusters (Figure 1E-G, Supplementary figure 2A), with clusters #3 and #5 predominantly derived from the TEB-containing sample (Figure 1F). To annotate these newly discovered fibroblast clusters, we created transcriptional scores for published fibroblasts from the developing fat pad^28,29^ and we found their enrichment in specific clusters in our dataset (Figure 1H). This comparative analysis with other single cell analyses allowed us to annotate clusters #0, #2, #4 as preadipocytes, adipocyte regulatory cells (AREGs) and interstitial progenitors, respectively (Figure 1I). Cells of cluster #3 and #5 were enriched in TEB sample and cells of cluster #5 expressed typical markers of proliferation (*Mki67*, *Top2a*, *Cdk1*, Figure 1G, Supplementary figure 2A) and were therefore annotated as peri-TEB and proliferative peri-TEB fibroblasts, respectively (Figure 1I). Finally, cells belonging to cluster #1 were predominantly derived from the ductal sample, were clearly distinct from fat pad fibroblasts but showed similarity to one fibroblast cluster previously identified in the adult mammary gland (Supplementary figure 2B)^26^; therefore, we annotated them as peri-ductal fibroblasts (Figure 1I).

We then used the transcriptional signatures of each cluster to inform on potential functional differences between these types of fibroblasts. We noticed that peri-TEB fibroblasts were highly enriched in contractility-related genes compared to the other clusters (Figure 1J) but expressed fewer genes encoding for fibrillar collagens and basal membrane proteins (Figure 1K). On the other hand, peri-TEB fibroblasts expressed tenascin C (Figure 1K), a gene encoding an anti-adhesive ECM molecule linked to organ development and cancer invasion^30,31^. Peri-TEB fibroblasts also differed in their expression of ECM remodeling enzymes, such as matrix metalloproteinases (*Mmps*), expressing more membrane-type *Mmps* (*Mmp14, Mmp15, Mmp16*, Supplementary figure 2C) that are important for mammary pubertal branching morphogenesis^32^, and fewer soluble *Mmps* (*Mmp2, Mmp3*). We next sought to computationally predict specific paracrine interactions and signaling networks between different fibroblast clusters and epithelial cells (both basal and luminal cells) using the algorithm CellChat, a bioinformatic tool designed to predict significant ligand-receptor interactions between two cell types from scRNAseq data^33^. Interestingly, this analysis identified peri-TEB fibroblasts as the strongest signaling hub, followed by preadipocytes and periductal fibroblasts (Supplementary figure 3A-C). Analysis of signals emanating from fibroblasts and received by epithelial cells revealed, surprisingly, that FGF ligands were not expressed by peri-TEB fibroblasts but rather by fat-pad associated fibroblasts (*Fgf2* by interstitial progenitors, *Fgf7* by AREGs and *Fgf10* by preadipocytes, Supplementary figure 3D-G). Peri-TEB fibroblasts, on the other hand, expressed higher levels of WNT ligands (*Wnt2, Wnt11*) and the WNT signaling regulator *Rspo1* (Supplementary figure 3D-G).

### A spatial atlas of mammary fibroblasts during pubertal branching morphogenesis

To retrieve a more precise information about the spatial distribution of the different fibroblast clusters identified in our scRNAseq dataset, we stained histological sections to analyze the expression of genes that we found to be characteristic of each fibroblast cluster based on the transcriptomic data (Figure 2A; Supplementary figure 4A) and annotated the regional localization of each examined field as containing predominantly ducts, TEBs or epithelium-free fat pad (Figure 2B). Expression of specific markers highly correlated with a defining regionalization of the different fibroblast clusters. Besides finding pan-fibroblastic markers (COL1A1, VIM) in all examined regions, we could detect expression of peri-TEB fibroblast markers (αSMA, NES, TNC, MYH10, SDC1) specifically in fibroblasts around the mammary TEBs, whereas the peri-ductal cluster marker *Mfap4* was found in fibroblasts lining the epithelial ducts (Figure 2C, D). Furthermore, genes enriched in fat pad-associated fibroblasts (CD34 for all of them, *F3* for AREGs, DPP4 and *Pi16* for interstitial progenitors, *Enpep* and *Thbs1* for preadipocytes) were indeed located in the distal fat pad devoid of mammary epithelium (Supplementary figure 4A-C). More specifically, AREGs defined by cluster #2 were identified lining blood vessels as expected from studies in the male fat pad^29^, preadipocytes were dispersed between adipocytes, and interstitial progenitors were found in the fat pad septae and in the interstitial reticulum (Supplementary figure 4D), a loose connective tissue at the fat pad border that creates a continuum with the septae dividing the fat pad lobules^28^.

**Fig. 2.**
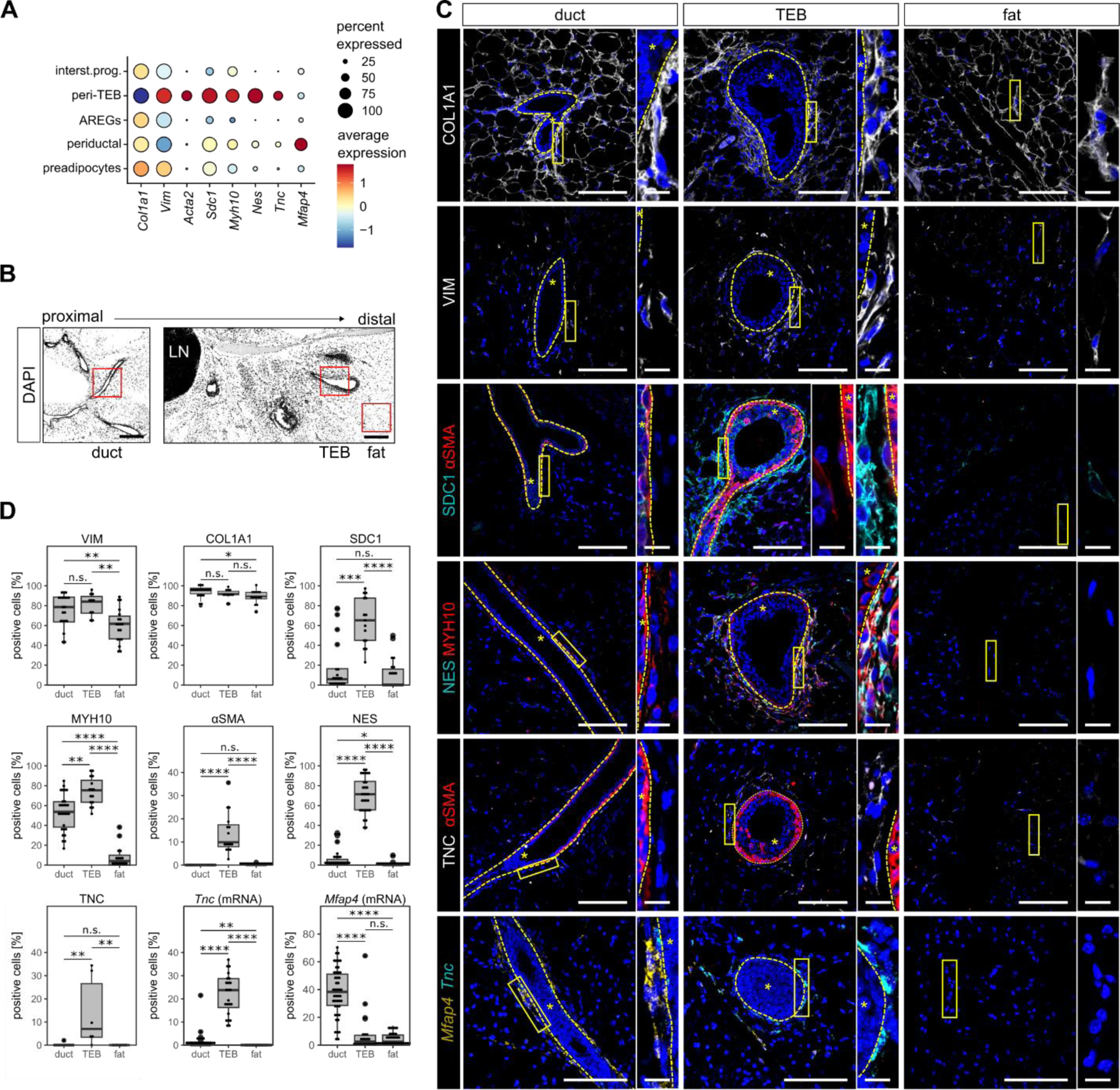
Spatial atlas of mammary fibroblasts during pubertal morphogenesis. **A.** A dot plot showing expression of marker genes used for spatial mapping. **B.** An overview of mammary gland on histological slide, red squares show fields of view (FOVs) containing duct, TEB or distal fat pad. Scale bar: 100 µm. **C.** Representative images showing expression of various gene or protein markers in regions containing duct, TEB or distal fat pad, the inset shows detail on fibroblasts. * marks epithelial compartment encircled by yellow dashed line. Scale bar: 100 µm, 10 µm in detail. **D.** Quantification of marker-positive stromal cells in different regions of mammary gland, shown as box plot. Each dot represents a single FOV, n = 3 biological replicates, N = 69 FOVs for VIM, 47 FOVs for COL1A1, 57 FOVs for SDC1, 59 FOVs for MYH10 and αSMA, 83 FOVs for NES, 26 FOVs for TNC, 113 FOVs for *Tnc* and *Mfap4*; statistical analysis: Wilcoxon test.

This spatial mapping demonstrated that the 6 fibroblast clusters that we identified by scRNAseq show a high degree of spatial organization within the mammary fat pad.

### The peri-epithelial fibroblasts expand during puberty

During pubertal growth, the mammary epithelium expands from a rudimentary tree to form a complex branched ductal system that entirely fills the fat pad^6,27^. To assess the dynamic behavior of the different fibroblast populations identified here during such prominent organ remodeling, we sought to sort different mammary fibroblasts by flow cytometry. To this aim, we first used a negative selection (Lin^neg^/Ep^neg^ [CD45^neg^/CD31^neg^/CD24^neg^/CD49f^neg^]) to exclude immune, endothelial and mammary epithelial cells^23,34,35^. Then, we designed a FACS gating strategy based on our newly identified markers distinguishing different types of mammary fibroblasts at the RNA level (Supplementary figure 5A) and by immunostaining (Supplementary figure 5B, C), dividing Lin^neg^/Ep^neg^ stromal cells in SCA1^neg^/CD39+ (encoded by *Entpd1*) peri-epithelial fibroblasts and SCA1+ (encoded by *Ly6a*) fat-pad associated fibroblasts. The latter ones were further separated into SCA1+/DPP4^neg^ cells comprising preadipocytes and AREGs, and SCA1+/DPP4+ interstitial progenitors (Figure 3A, Supplementary figure 5D, E). Immunofluorescence staining of freshly isolated FACS-sorted cells from pubertal mammary glands confirmed the presence of αSMA+ and NES+ peri-TEB fibroblasts in the SCA1^neg^/CD39+ population (Supplementary figure 5F-I). Flow cytometry analysis of mammary glands at 3 weeks of age (pre-puberty), 5 weeks (peak of puberty) and 14 weeks (adult stage) revealed the expansion of peri-epithelial SCA1^neg^/CD39+ fibroblasts relatively to other fibroblast populations (Figure 3B, C), consistent with the prominent increase in epithelial cells due to ductal extension and ramification. In addition, our transcriptomic map hinted to the presence of proliferative fibroblasts in TEB proximity (Figure 1F, G) and indeed a 2-hour EdU pulse revealed EdU+ stromal cells mostly located near TEBs (Supplementary figure 6A, B). Intriguingly, we detected proliferative EdU+/SDC1+/αSMA+ fibroblasts in the peri-TEB niche (Supplementary figure 6C).

**Fig. 3.**
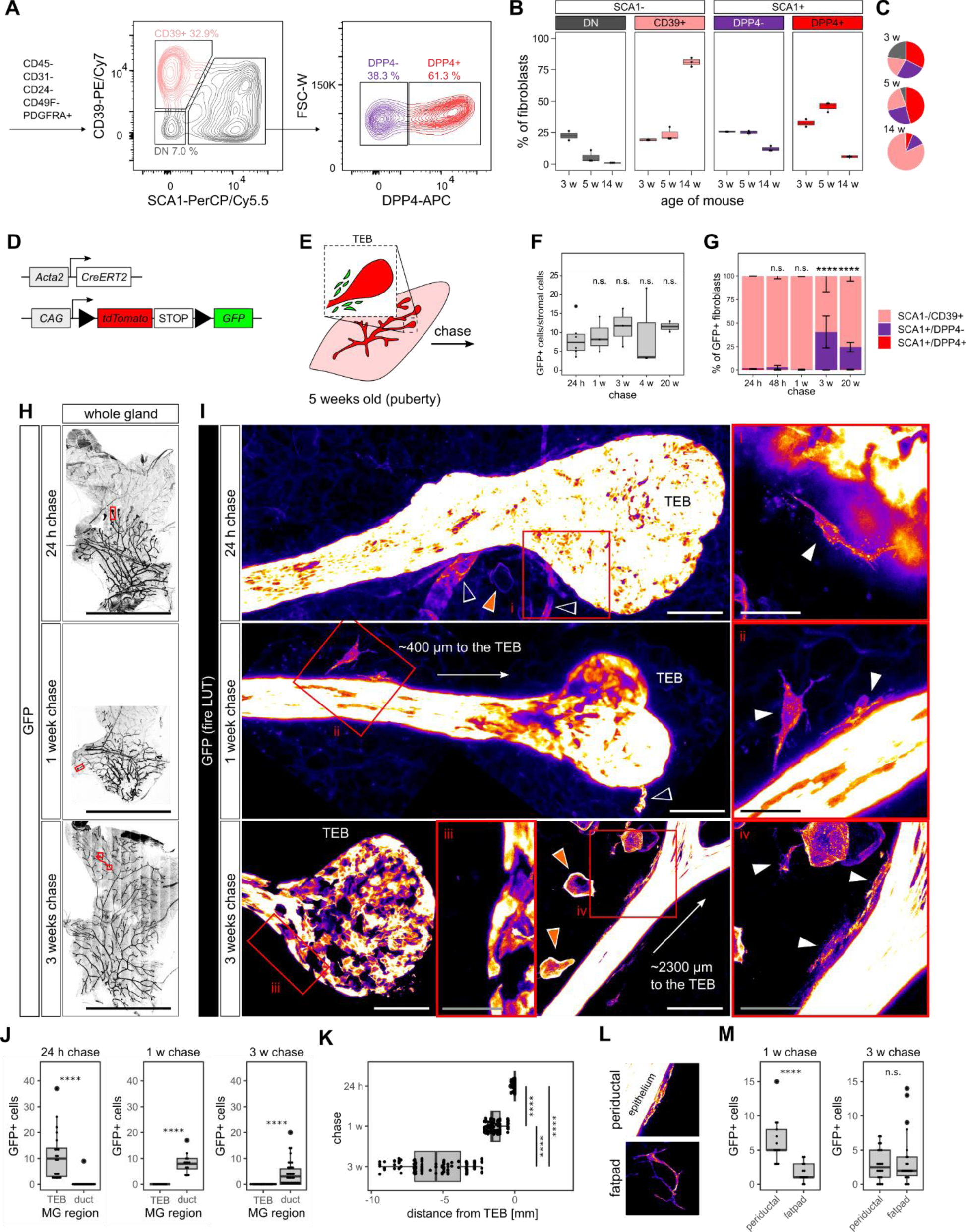
Peri-TEB fibroblasts are a transient cell state and do not migrate with the invading epithelium. **A.** Representative FACS plots dividing mammary fibroblasts into CD39+; SCA1+/DPP4- and SCA1+/DPP4+ populations. **B, C.** Flow cytometry quantification of mammary fibroblast proportions from 3 week, 5 week and 14 week old mice shown as boxplots (**B**) and pie charts (**C**), n = 3 mice per sample. **D.** Schematic representation of the *Acta2-CreERT2;R26-mT/mG* mouse model. **E.** Induction strategy for the chase experiments. **F, G.** Flow cytometry quantification of GFP+ cells out of total stromal cells (**F**) and quantification of the proportion of FACS populations within the GFP+ fibroblasts (**G**); n = 3 mice per time-point, statistical analysis: Wilcoxon (**F**) and chi square test (**G**). **H, I.** Projections of z-stack imaging of whole mammary glands after tamoxifen induction and 24 h-, 1 week- or 3 week-long chase. Red boxes in whole organ overview pictures (**H**) indicate regions presented in **I.** Red boxes in detail pictures of ductal or peri-TEB regions (**I**) indicate magnified regions with GFP+ fibroblasts (i-iv). GFP channel is shown as “fire” lookup table. White arrowheads indicate GFP+ fibroblasts, orange arrowheads indicate GFP+ adipocytes, empty arrowheads indicate GFP+ mural cells. Scale bars: 1 cm in **H**; 100 µm in **I**, 50 µm in detail in **I**. **J.** Quantification of positional distribution of GFP+ fibroblasts after 24 h or 3 weeks of chase; n = 3 mice, N = 38 z-stack for the 24 h chase, 22 z-stacks for 1 week and 51 z-stacks for the 3 weeks chase, statistical analysis: Wilcoxon test. **K.** Quantification of distance of GFP+ fibroblasts from the TEB; n = 3 mice, N = 343 fibroblasts for 24 h, 106 for 1 week and 144 for 3 week-long chase. **L, M.** Representative appearance (**L**) and quantification (**M**) of GFP+ fibroblasts found after 1 or3 weeks of chase. n = 3 mice; N = 22 z-stacks for 1 week and 51 z-stacks for 3 weeks chase (M).

These results indicate that, at puberty, peri-epithelial fibroblasts expand alongside the growing ductal epithelium and that proliferating fibroblasts are specifically concentrated around TEBs.

### Peri-TEB fibroblasts are trailing behind the growing epithelium

To assess the fate of the peri-TEB fibroblasts and understand whether they contribute to the expansion of peri-epithelial fibroblasts, we used a genetic labeling approach to lineage trace them and to follow their behavior *in vivo*. For this, *Acta2-CreERT2;R26-mT/mG* mice were induced at the peak of puberty to label peri-TEB fibroblasts with permanent and heritable GFP expression (Figure 3D-E). After 24/48 hours (acute labeling) or longer chase times (1, 3, 4, 20 weeks), we analyzed the GFP+ fibroblasts by flow cytometry. Surprisingly, flow cytometry showed no expansion of GFP+ cells within the stromal cell population (Figure 3F). However, the distribution of GFP+ cells in the fibroblast populations considerably changed: while acute labeling and 1 week chase resulted in GFP+ cells only in the peri-epithelial CD39+ population, longer chase times for 3 weeks or 20 weeks resulted in a substantial increase in SCA1+/DPP4^neg^ cells (in purple in Figure 3G, Supplementary figure 7A), suggesting that *Acta2*+ peri-TEB fibroblasts can give rise to preadipocytes.

To visualize the spatial distribution of *Acta2*+ peri-TEB fibroblasts and their progeny, we employed whole-organ imaging of CUBIC-cleared mammary glands^36,37^, a method that enables evaluation of a large volume of the organ to facilitate the detection of rare cells or cells with an unconventional 3D shape, like fibroblasts. With a short pulse of labeling, the GFP+ fibroblasts were localized around TEBs, in contact with the epithelial cells (Figure 3H-J; note that the level of GFP expression is considerably lower in fibroblasts than in the epithelial cells and mural cells [Supplementary figure 7B], therefore the epithelial signal is overexposed) whereas we could not detect any GFP+ fibroblasts surrounding the ducts (Figure 3J, Supplementary figure 7C). Immunostaining confirmed that the GFP+ cells found in the peri-TEB stromal region were indeed αSMA+/VIM+ and CD34^neg^/DPP4^ne^ peri-TEB fibroblasts (Supplementary figure 8A, B).

After 1 or 3 weeks of chase, however, there were no GFP+ fibroblasts around TEBs anymore and the *Acta2*-labeled fibroblasts appeared to have moved to the subtending epithelial ducts (Figure 3H-J, Supplementary figure 8C), at a distance from the TEBs ranging from 1 to 2 mm after 1 week chase and 2.5 mm to 10 mm after 3 weeks chase (Figure 3K), roughly corresponding to the distance covered by growing TEBs in 1 or 3 weeks^7^. Of interest, after a 1- or 3-week chase, we found GFP+ fibroblasts either sitting right on the mammary epithelial ducts or dispersed in the fat pad (Figure 3L, M) and by immunostaining we demonstrated that the GFP+ fibroblasts (VIM+) located either in the peri ductal stroma or further in the fat pad did not express αSMA (Supplementary figure 8C). This corroborates the flow cytometry results suggesting that both periductal fibroblasts and preadipocytes can arise from *Acta2*+ fibroblasts. Altogether, by lineage tracing analysis we provided evidence indicating that the peri-TEB fibroblasts do not prominently expand during puberty and do not seem to move forward to escort the invading epithelial TEBs, but they rather lag behind the growing ducts and either differentiate or give rise to peri-ductal fibroblasts and later to preadipocytes.

### Peri-TEB fibroblasts are recruited from the fat pad stroma by the growing epithelium

The lineage tracing experiments described above drew the important conclusion that the peri-TEB fibroblasts are left behind during the ductal growth, raising the question of the origin of new contractile fibroblasts that must be delivered to TEBs to maintain an equilibrium within the peri-TEB stromal cells and to support epithelial growth and fat pad invasion. We reasoned that the tips of growing TEBs inevitably encounter preadipocytes dispersed within the fat pad on their morphogenetic journey, therefore we aimed to lineage trace preadipocytes to establish whether they can be recruited by growing TEBs and acquire a peri-TEB fate. We observed that the Notch1 receptor was expressed by preadipocytes, although not exclusively since it is also found in peri-epithelial fibroblasts (Figure 4A). We therefore used pubertal *Notch1-CreERT2;R26-mT/mG* mice for lineage tracing (Figure 4B, C) and quantified and visualized the distribution of labelled fibroblasts by flow cytometry and microscopy. Similar to *Acta2-CreERT2* tracing, *Notch1*-expressing stromal cells did not expand (Figure 4D), but the proportion of labelled fibroblasts in different subsets changed. Upon acute labeling (24 h), most GFP+ fibroblasts were found in the SCA1+/DPP4^neg^ gate (Figure 4E), suggesting an efficient labeling of preadipocytes; however, already after 1 week of chase, the ratio switched, and we detected GFP+ fibroblasts primarily within the SCA1^neg^/CD39+ population of peri-epithelial fibroblasts (Figure 4E). Consistently, by whole-tissue microscopy, we found GFP+ fibroblasts both dispersed within the fat pad and touching the epithelium at acute labeling, whereas 1 week after labeling the GFP+ cells either formed clusters of adipocytes in the fat pad or they were located around TEBs (Supplementary figure 9).

**Fig. 4.**
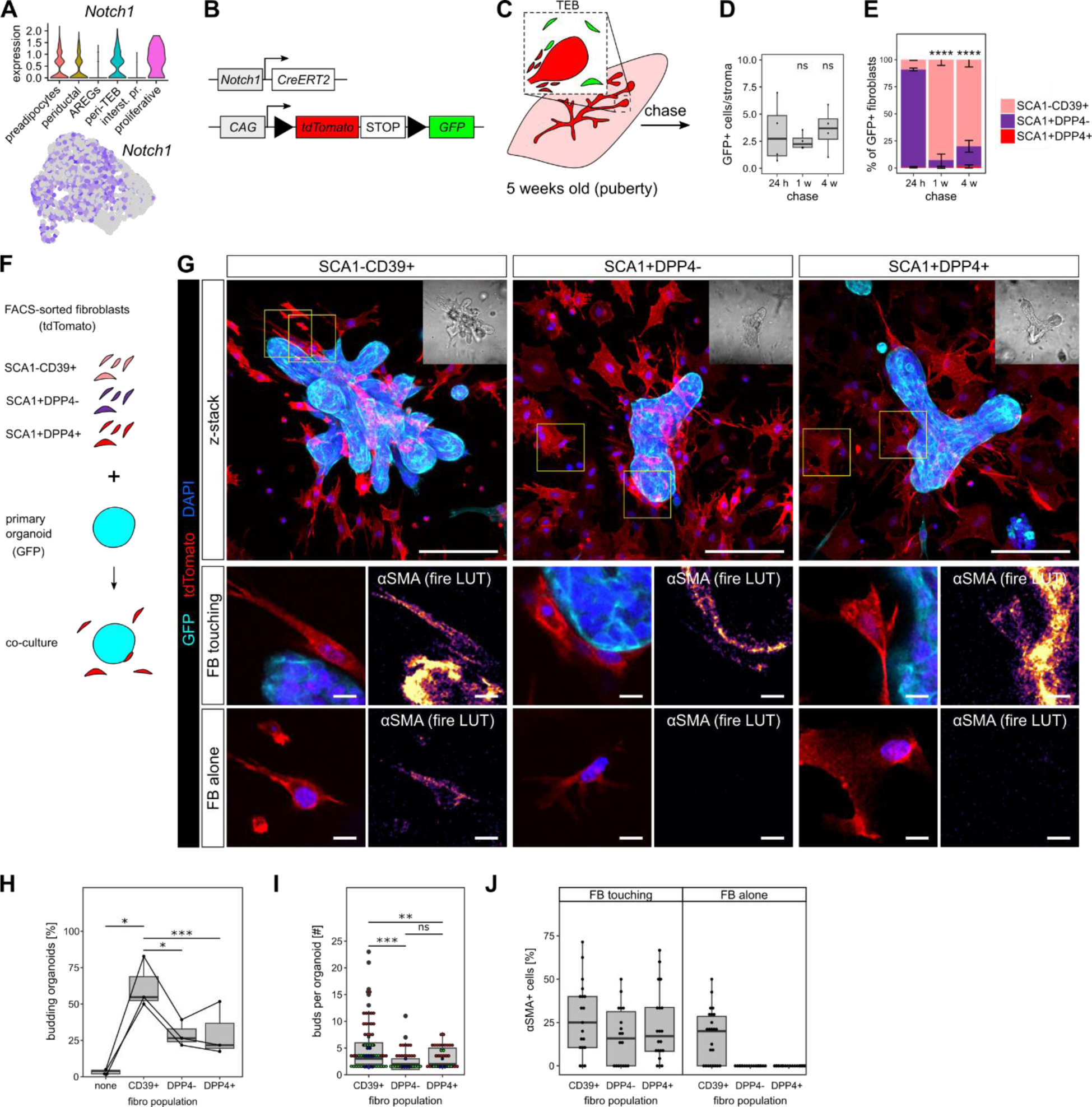
Growing epithelium recruits peri-TEB fibroblasts from the fat pad. **A-E.** *Notch1* fibroblast chase experiment. **A.** Violin plot and UMAP expression of *Notch1* in fibroblast clusters. **B.** Schematic representation of the *Notch1-CreERT2;R26-mT/mG* mouse model. **C.** The induction strategy. Flow cytometry quantification of GFP+ cells out of total stromal cells (**D**) and quantification of the proportion of FACS populations within the GFP+ fibroblasts (**E**); n = 3 mice per time-point, statistical analysis: Wilcoxon (D) and chi square test (**E**). **F-J.** Co-culture of sorted fibroblasts with epithelial organoids. **F.** A schematic representation of the experiment. **G.** Representative images of the organoids after 5 days of co-culture with FACS-sorted fibroblasts (organoids in cyan, fibroblasts in red). The inlets show brightfield images of the cultures. The magnified regions (demarcated by yellow lines) provide a detailed view on fibroblasts in contact with the organoid or further away from it in the ECM and their staining for αSMA. Scale bars: 100 µm and 10 µm in detail. **H.** Quantification of percentage of organoid budding, shown as box plot. Each dot represents a biological replicate, lines connect paired experiments; n = 3 independent experiments, statistical analysis: Student’s t-test. **I.** Quantification of the number of buds formed per organoid, each dot represents one organoid, colors code paired experiments; n = 3 independent experiments, N = 91, 40 and 44 organoids for CD39+, DPP4- and DPP4+ population, respectively, statistical analysis: Student’s t-test. **J.** Quantification of αSMA+ fibroblasts (FB) in co-cultures, shown as a box plot, each dot shows one organoid; n = 3 independent experiments, N = 24, 18 and 20 organoids for CD39+, DPP4- and DPP4+ population, respectively.

The switch in *Notch1-*traced cells from SCA1+/DPP4^neg^ to SCA1^neg^/CD39+ cell subsets may suggest that preadipocytes change state by becoming “activated” and convert into contractile peri-epithelial fibroblasts when they encounter a growing duct. We cannot however exclude the possibility that the small population of SCA1^neg^/CD39+ cells labeled at 24 h expanded and generated the GFP+ peri-epithelial fibroblasts that we observed at later time points after tracing.

In an attempt to discriminate between these two possibilities, we conducted a transplantation experiment, engrafting tdTomato+ fragments of mammary glands (consisting of epithelium and surrounding stroma both labeled by red fluorescence) into cleared fat pads of immunocompromised host mice (Supplementary figure 10A). Supporting our hypothesis that fibroblasts can be recruited by growing TEBs and they do not migrate alongside the growing mammary tips, 3.5 weeks after transplantation, the tdTomato+ epithelium reconstituted a branched outgrowth (Supplementary figure 10B) presenting red fluorescent TEBs, that were surrounded by tdTomato-negative host-derived stromal cells, while tdTomato+ engrafted stromal cells remained at the original grafting site, corroborating our lineage tracing results (Supplementary figure 10C, D).

Furthermore, we co-cultured GFP+ mammary epithelial organoids with the different subsets of tdTomato-labeled fibroblasts that we can now sort by flow cytometry (SCA1^neg^/CD39+, SCA1+/DPP4^neg^ and SCA1+/DPP4+; Figure 4F, G). In such co-cultures, we have recently shown that fibroblasts induce organoid branching in a contractility-dependent manner^17^. Surprisingly, we found that after 5 days of co-culture, fibroblasts of all three populations promoted epithelial organoid budding, although the peri-epithelial SCA1^neg^/CD39+ fibroblasts displayed the highest morphogenetic potential (Figure 4G-I). Therefore, we probed αSMA expression in fibroblasts co-cultured for 5 days with mammary epithelial organoids and discovered that all fibroblast populations acquired αSMA expression when in direct contact with epithelial organoids (Figure 4G, J). Together these results demonstrate that the growing epithelium can recruit fibroblasts from the fat pad and induce their peri-TEB phenotype and namely contractility.

## Discussion

### Heterogeneity of mammary fibroblasts

Dissection of fibroblast heterogeneity is a blooming topic of research, thanks to scRNAseq analyses that can now offer unprecedented details into the differences between individual fibroblast subtypes presenting common features as well as organ specific differences^4,5^. In the mammary gland, our recent scRNAseq analyses discriminated two distinct clusters in the embryonic mammary stroma^38^; also, other groups identified four fibroblast clusters in the adult mammary gland^26^, revealed changes in fibroblast abundance and ECM expression in the aging mammary glands^25^ and compared the profile of normal fibroblasts with breast cancer-associated fibroblasts^23^. We have recently reported the existence of specialized αSMA+ peri-TEB fibroblasts that exist only in the proliferating and branching pubertal mammary gland^17^. In the present study, we employed our newly discovered cluster-specific markers to draw a full picture of the spatial distribution of different fibroblasts within the mammary stroma (Figure 5). We show that three SCA1+ clusters represent fat pad fibroblasts, previously found in the male fat pad^28,29^ as well as in the adult mammary gland^26^. In this study, we focused on two clusters of peri-epithelial fibroblasts that reside in the collagen-rich sheath around the epithelium: periductal and contractile peri-TEB fibroblasts, unique to puberty.

**Fig. 5.**
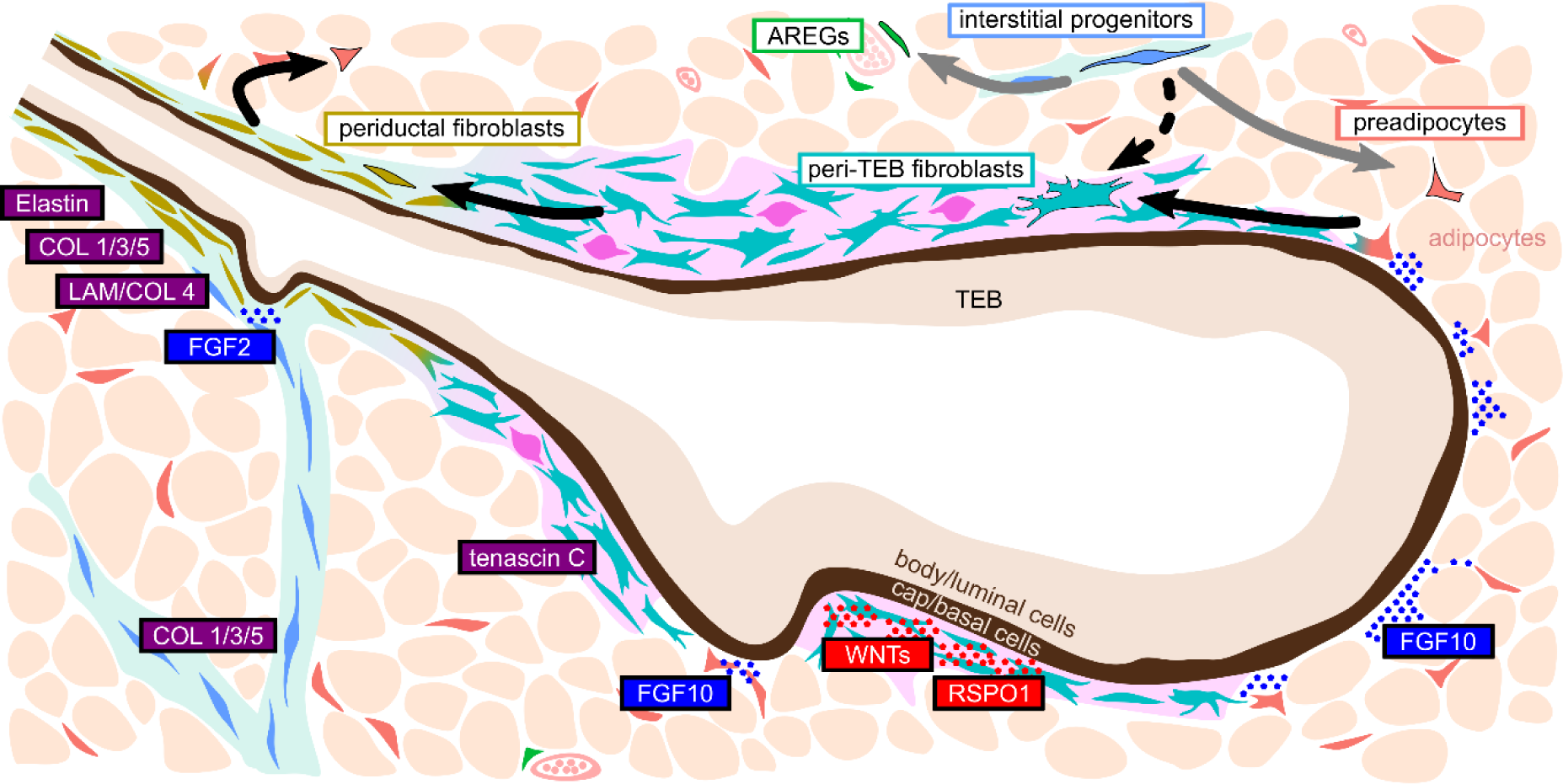
A scheme: contractile peri-TEB fibroblasts are recruited by the growing epithelium. A schematic representation of our findings. Mammary stroma comprises 5 clusters of fibroblasts (+ proliferative fibroblasts) with distinct spatial locations: peri-TEB fibroblasts surround TEBs, periductal fibroblasts wrap around subtending ducts, interstitial progenitors reside in collagenous fat pad septae, AREGs dwell in blood vessel stroma, and preadipocytes are scattered between adipocytes. Interstitial progenitors have been shown to provide other fat pad fibroblasts (Stefkovic et al. 2021, gray arrows). Our data suggest that fat pad fibroblasts are recruited by the TEB and activated into peri-TEB phenotype, then differentiate into periductal fibroblasts and back to preadipocytes (black arrows). Based on spatial resolution of fibroblast clusters, we predict spatial resolution of ECM (purple), WNT signaling ligands (red) in peri-TEB stroma and FGF signaling ligands (blue) in fat pad and fat pad septae. The dots show expected sites where mammary epithelium encounters FGF/WNT ligands.

Our data thus suggest a strong correlation between spatial organization of stromal cells and fibroblast heterogeneity, that seems to be conserved in other organs. For example in the skin, papillary fibroblasts are close to the epidermis, while SCA1+ reticular fibroblasts are embedded deeper in the dermis, close to the subcutaneous fat pad^39^; likewise, in the liver, SCA1+ periportal fibroblasts sit in the portal area and form a separate scRNAseq cluster, distinguished from hepatic stellate cells that line hepatocytes^40,41^. We thus propose that the spatial patterning of fibroblast subtypes may reflect the niche-specific needs for ECM as ECM-related genes represent the major discriminators of fibroblast subtypes across different organs^5^.

### Fibroblast hierarchy

Fibroblasts are stromal cells of remarkable plasticity, that has been revealed by genetic lineage tracing and transplantation assays, helping to dissect cellular hierarchies in various tissues. In the skin, papillary and reticular fibroblast lineages split during embryonic development and never overlap in adult homeostasis^39^. In the fat pad, Dpp4+ interstitial progenitors contribute to both preadipocytes and AREGs in a unidirectional way^28,42^ in accord with transcriptomic analysis that identified Pi16+/Dpp4+ fibroblasts as a progenitor population sitting at the top of the fibroblast hierarchy^4^. Our lineage tracing experiments show that the newly discovered peri-TEB fibroblasts can contribute to both periductal fibroblasts and preadipocytes, but in accord with aforementioned studies, we never observed them to give rise to Pi16+/Dpp4+ interstitial progenitors. Importantly, we show by transplantation and ex vivo recombination that both preadipocytes and interstitial progenitors can be activated by contact with the epithelium and acquire a peri-TEB contractile phenotype. Thus, our results support the hypothesis that peri-TEB fibroblasts represent a transient cell state associated with epithelial morphogenesis and provide a link between fat pad fibroblasts and peri-epithelial fibroblasts, demonstrating that they are not separate lineages (Figure 5).

Fibroblasts can also differentiate into pericytes and adipocytes in the skin^43,44^ and in the interscapular, inguinal^45^, mesenteric and gonadal fat pad^42^. In the mammary gland, adipocytes dedifferentiate into fibroblasts during pregnancy and are reestablished from fibroblasts during involution^46,47^. It is yet to be determined which fibroblast clusters can trans-differentiate into adipocytes in homeostasis and during pregnancy-associated changes of the mammary stroma. Our data showing the remarkable plasticity of these cells, at least in co-cultures, suggest however that both fat pad and peri-epithelial fibroblasts could be able to give rise to adipocytes.

### Fibroblasts orchestrate epithelial morphogenesis

Fibroblasts are believed to be critical regulators of mammary branching morphogenesis^2,10^, due to their capacity of secreting Fibroblast Growth Factors (FGFs)^11^. FGF10 acts as a chemoattractant in organoids^11^ and FGF10-soaked beads locally accelerate branching when implanted *in vivo*^8^. Our transcriptomic analysis shows that FGFs are mostly expressed by fat pad fibroblasts (Figure 5). While most epithelial cells are shielded from the fat pad by a dense collagenous stroma, the leading edge of TEB is in direct contact with FGF10-expressing preadipocytes. Thus, we believe that the spatial organization of fibroblast clusters can explain how FGF10-mediated chemoattraction acts *in vivo*. Also FGF2 induces epithelial proliferation and organoid budding *in vitro*^12^, which is reminiscent of mammary side branching^48^. Here we found that FGF2 is produced by interstitial progenitors, located along the septae dividing the fat pad; these stromal cells could stochastically encounter a growing duct however if this impacts the pattern of epithelial branching is an attractive hypothesis that remains to be mechanistically explored. We have also detected cluster-specific Wnt ligand expression (Figure 5), with peri-TEB fibroblasts being the stronger producers of Wnt ligands, suggesting that they may create a stem cell-promoting niche, rich in Wnt ligands, around the TEBs. Other paracrine signals predicted to act between fibroblasts and epithelium by the Cellchat analysis we provide here remain to be functionally addressed.

ECM expression and organization is another critical regulator of mammary epithelial branching^49–51^ and our scRNAseq analysis reveals distinct patterns of ECM expression in different fibroblast clusters. Our analysis shows that peri-TEB fibroblasts express low levels of fibrillar collagens, including *Col1a1* and *Col1a2*, suggesting that they may produce the collagen sheath only after their transition into peri-ductal fibroblasts. Similarly, we detected lower expression of basal membrane genes in peri-TEB fibroblasts, in agreement with the well documented reduction in basal membrane thickness around TEBs^27^. Peri-TEB fibroblasts are also the primary source of membrane bound Matrix Metalloproteases MMPs that can further promote basal membrane thinning and have been shown to facilitate mammary branching *in vivo*^32^. Finally, the peri-TEB fibroblasts we characterize here are the exclusive producers of Tenascin C, another ECM molecule associated with pro-invasive properties^30,31^.

A new line of evidence suggest that stromal cells may also regulate epithelial morphogenesis through mechanical forces derived from actomyosin cell contraction, as it was shown for the regulation of lung branching^18,52^, the growth of feathers^53^ and hair follicles^54^ as well as gut vilification^55^. In the mammary gland, we have recently demonstrated that fibroblast contractility drives epithelial organoid branching in a co-culture system^17^. It is conceivable that the contractile fibroblasts positioned around the neck of TEBs that we studied here may create a hoop stress that facilitates the ductal invasion through the fat pad, as previously suggested for myoepithelial cells^56^. Of note, the peculiar localization of contractile peri-TEB fibroblasts correlates with the transition of the cuboidal epithelial cap cells of TEBs into myoepithelial cells lining the subtending ducts, opening an exciting opportunity to test the involvement of contractile fibroblasts in this process in further studies. In conclusion, we have characterized a novel fibroblast population that creates a specialized mechano-chemical niche to facilitate epithelial morphogenesis (Figure 5).

### Fibroblasts activation – signals from the epithelium and beyond

The process of fibroblast activation is probably best studied in wound healing, when fibroblast contractility and ECM production is turned on to aid proper wound closure and healing. Excessive fibroblast activation, however, can result in scarring, fibrosis and provide cancer prone microenvironment. Contractile fibroblasts, however, are also found in developing tissues and it is not clear if they are controlled by conserved mechanisms and molecular regulators. Interestingly, we have observed that while peri-TEB fibroblasts are characterized by a signature of contractility, they express lower levels of fibrillar collagens, suggesting that the activation process is either completely different or only partial, when compared to wound healing. Many of the signaling pathways commonly associated with fibroblast activation have been shown to be involved in mammary fibroblasts regulation, like PDGF^57^, FGF^58^ or WNT signaling^59–61^. Finally, contact with macrophages can activate myofibroblasts in wounds, via cell-cell adhesion and TGFβ signaling^62^, while macrophages are present in the peri-TEB stroma and their depletion impairs mammary outgrowth^63,64^. It is therefore tempting to speculate that TEBs invading the fat pad create a miniature wound-like environment, involving activation of contractile fibroblasts that are recruited to propel epithelial branching in an evolutionary conserved manner. Further research into fibroblast activation during organ development is needed to understand the differences and similarities between morphogenesis, wound-healing, fibrosis and cancer.

## Material and methods

### Mice

All procedures involving animals were performed under the approval of the Ministry of Education, Youth and Sports of the Czech Republic (license # MSMT-9232/2020-2), supervised by the Expert Committee for Laboratory Animal Welfare of the Faculty of Medicine, Masaryk University, at the Laboratory Animal Breeding and Experimental Facility of the Faculty of Medicine, Masaryk University (facility license #58013/2017-MZE-17214), or under the approval of the ethics committee of the Institut Curie and the French Ministry of Research (reference #34364-202112151422480) in the Animal Facility of Institut Curie (facility license #C75–05–18). ICR mice and nude mice were obtained from the Laboratory Animal Breeding and Experimental Facility of the Faculty of Medicine, Masaryk University. *LifeAct-GFP* mice were created by Wedlich-Söldner team^65^, *R26-mT/mG*^66^ and *Acta2-CreERT2* mice^67^ were acquired from the Jackson Laboratory. *Notch1-CreERT2* mice^68^ were created in our laboratory. Transgenic animals were maintained on a C57BL/6 background. Experimental animals were obtained by breeding of the parental strains, the genotypes were determined by genotyping. The mice were housed in individually ventilated or open cages, all with ambient temperature of 22°C, a 12 h:12 h light:dark cycle, and food and water ad libitum. For induction of *CreERT2* constructs, mice were treated with tamoxifen (Sigma, 2 mg/10 g of mouse) diluted in corn oil (Sigma), delivered via intraperitoneal injection. For EdU incorporation assay, 5 weeks old wild-type females were injected with 10 nmol/1 g of mouse, delivered via intraperitoneal injection 2 h prior to euthanasia. Mice were euthanized by cervical dislocation and mammary gland tissues were collected immediately.

### Microdissection of TEB/duct

For microdissection of TEB- and duct-containing regions, 5 weeks old *R26-mT/mG* females were used. Mammary glands #3 and #4 were dissected and spread on 10 cm plastic petri dishes. Under the control of a fluorescent stereoscope (Leica FM165C), 0.5×0.5 mm pieces of tissue containing either TEBs or pieces of ducts (assessed based on tdTomato fluorescence) were dissected and collected in CO_2_-independent medium on ice. 5-8 mice were pooled for one experiment.

### Single cell RNA sequencing

For scRNAseq experiment, microdissected pieces of mammary gland were digested in a collagenase/hyaluronidase solution [3 mg/ml collagenase A, 100 U/ml hyaluronidase (all Merck), 5% fetal bovine serum (FBS; Hyclone/GE Healthcare) in CO_2_ independent medium (Thermo Fisher Scientific)] for 2 hours at 37°C, shaking at 120 rpm. Resulting tissue fragments were treated with 0.25% trypsin, 5 mg/ml dispase II, 100 µg/ml DNase and red blood cell lysis buffer (Sigma) and filtered through a 40 µm cell strainer. Resulting single cell suspension was pooled down, resuspended in freezing medium (10% DMSO in FBS) and frozen by slow freezing at −80°C and moved to liquid nitrogen 24 h thereafter. Sequencing was performed at Single Cell Discoveries BV, Netherlands, using the 10x Genomics platform, at 5,000 target cells/sample and with sequencing depth of 50,000 reads per cell.

### scRNAseq data analysis – data preprocessing

The analysis of scRNAseq data was performed in R (R systems) using the Seurat package^69^. The following filters were applied to the data to remove doublets and damaged cells: genes with expression in < 3, *nCount_RNA* < 3000 or > 60000, *nFeature_RNA* < 1000 or with a percentage of mitochondrial reads > 20 were removed. The data were normalized with the *SCTransform* function. Uniform Manifold Approximation and Projection and neighbors identification were computed using the *RunUMAP* and *FindNeighbors* function on the first 9 PCs Clusters were defined using the *FindClusters* function with a resolution set to 0.2. Fibroblast, epithelial and leukocyte populations were then identified based on canonical markers (*Col1a1*, *Col3a1*, *Pdgfra* for fibroblasts; *Ptprc* for leukocytes; *Epcam*, *Cdh1* for epithelium).

### scRNAseq data analysis – subpopulation analysis

The fibroblast, epithelial and leukocyte subsets of cells were then merged together and reanalyzed using the first 10 PCs for *RunUMAP* and 0.3 resolution for *FindClusters.* Possible lymphocyte-containing doublets within fibroblast cluster were identified by *Ptprc* expression and excluded from the analysis. Cluster markers were calculated using the *FindAllMarkers* function. The expression scores were calculated using the *AddModuleScore* function with genes in supplementary table 1.

### scRNAseq data analysis – cell communication analysis

Communication link between cell populations were identified using the CellChat R package^33^. Only epithelial and fibroblast clusters were used for this analysis. Interactions between the different clusters were identified based on the mouse ‘Secreted Signaling’ internal CellChat database. Interactions identified in less than 5 cells per group were filtered out. Interactions between groups were visualized using the ‘netVisual_chord_gene’ function.

### Immunohistochemistry (IHC)

For IHC, mammary glands #4 were dissected, spread on glass slide and fixed in 4% paraformaldehyde (EM grade; Electron Microscopy Sciences) overnight at 4°C, dehydrated in ethanol solutions with increasing concentration and xylene and embedded in paraffin. 5 µm sections were cut on rotational microtome (Thermo scientific, Microm HM340E). After rehydration, sections were boiled in pH6 citrate buffer or pH9 Tris-EDTA buffer to retrieve antigens, blocked in 1% bovine serum albumin and 5% FBS and incubated with primary antibodies (supplementary table 2), fluorophore-conjugated secondary antibodies (ThermoFisher Scientific) and 1 μg/ml 4′,6-diamidino-2-phenylindole (DAPI, Merck), mounted (Aqua Poly/Mount, Polysciences) and imaged on laser scanning confocal microscope (LSM780/800/880/900, Zeiss). For SCA1 staining, slides were first stained for COL1A1 than biotin-conjugated SCA1 primary antibody was followed by incubation with 1 µg/ml Alexafluor633-conjugated streptavidin (ThermoFisher Scientific) for 1 hour before counterstaining with DAPI. EdU detection was performed using Click-iT imaging kit (ThermoFisher Scientific) following the manufacturer’s instructions. The quantification of positive cells was done manually in ImageJ, using DAPI signal to count total stromal cell number. Cells of epithelial ducts and blood vessels, recognized by typical morphology, autofluorescence and cell shape, were omitted from analysis. The field of views (FOVs) were all of the same size (212×212 µm) and were scored “duct”, “TEB” or “fat” based on morphology and position of the structures (TEBs: stratified epithelium, bulb-like shape and position of the structure at the distal part of the mammary epithelium (invasive front); duct: bi-layered epithelium; fat: no epithelial structures in the FOV).

### In situ hybridization (ISH)

For ISH, mammary glands were processed in the same way as for IHC and the staining was performed using RNA Scope kit following manufacturer’s instructions, using commercially available probes (supplementary table 2). The quantification of positive cells was done manually in ImageJ similarly to IHC quantification, using DAPI signal to count total stromal cell number. Rules for determining positive cell were unique for each probe, based on the level of expression (3 dots for *Enpep*, *Thbs1* and *Ly6a*; cytoplasm full of dots for *F3*, *Tnc, Pi16* and *Mfap4*).

### Fluorescence assisted cell sorting (FACS)

For FACS or flow cytometry, harvested mammary glands #1, #2, #3, #4 and #5 were chopped into 0.5×0.5×0.5 mm^3^ pieces and digested in a solution of collagenase and trypsin [2 mg/ml collagenase A, 2 mg/ml trypsin, 5 μg/ml insulin, 50 μg/ml gentamicin (all Merck), 5% FBS (Hyclone/GE Healthcare) in DMEM/F12 (Thermo Fisher Scientific)] for 30-60 min at 37°C. Resulting tissue suspension was treated with DNase I (20 U/ml; Merck) and red blood cell lysis buffer and filtered through 40 µm cell strainer to achieve single cell solution. The cells were resuspended in staining medium (10% FBS, 100 U/ml of penicillin, and 100 μg/ml of streptomycin in CO_2_ independent medium) with antibodies (supplementary table 2) and incubated on ice in dark for 30 min. Then, the cells were washed twice with PBS, resuspended in flow buffer (5 mM EDTA, 1% BSA, 1% FBS in phenol red-free DMEM/F12 with HEPES) and passed through a 40 µm cell strainer. The analysis was done on Aria III or Fusion (both BD), operated by SORP software (BD). The flow cytometry data were analyzed using the FlowJo software (BD).

### Immunocytochemistry (ICC)

For immunofluorescent analysis, 10,000 FACS-sorted fibroblasts were plated directly on coverslip-bottom 8 well plates (IBIDI) and cultured in fibroblast medium [10% FBS, 1× ITS (10 μg/ml insulin, 5.5 μg/ml transferrin, 6.7 ng/ml sodium selenite), 100 U/ml of penicillin, and 100 μg/ml of streptomycin in DMEM], fixed with 4% formaldehyde for 15 min, permeabilized with 0.5% Triton X-100 in PBS for 10 min and blocked with PBS with 10% FBS for 30 min. Then the cells were incubated with primary antibodies (supplementary table 2) for 2 h at RT. After washing, the cells were incubated with fluorophore-conjugated secondary antibodies (ThermoFisher Scientific) and phalloidin AlexaFluor 488 and DAPI for 2 h at RT. The cells were photographed using LSM 780/880/900 (Zeiss).

### Whole-mount clearing and imaging

Staining and clearing of the mammary glands was done following clear, unobstructed brain imaging cocktails (CUBIC) protocol^36,37^. Briefly, mammary glands #3 or #4 were harvested and fixed in in 4% paraformaldehyde overnight at 4°C, washed and incubated in CUBIC reagent 1 (25% (w/w) urea, 25% (w/w) N,N,N’,N’-tetrakis(2-hydroxypropyl)ethylenediamine, 15% (w/w) Triton X-100 in distilled water) for 4 days shaking at RT. After washing, the tissue was blocked using blocking buffer (5% FBS, 2% BSA, 1% Triton X-100, 0.02% sodium azide in PBS) overnight at RT, incubated with primary antibodies (supplementary table 2) diluted in blocking buffer for 3 days at 4°C with rocking, washed three times for 2 h (0.05% Tween 20 in PBS) and incubated with DAPI (1 µg/ml) in blocking buffer. Then the glands were transferred to CUBIC reagent 2 (50% (w/w) sucrose, 25% (w/w) urea, 10% (w/w) 2,2’,2’’-nitrilotriethanol, 0.1% (w/w) Triton X-100 in distilled water) for >1 day at RT with rocking. The samples were mounted with CUBIC reagent 2 between two coverslips with double-sided tape as a spacer to enable imaging from both sides and they were imaged on laser scanning confocal microscope LSM780/880 (Zeiss). Non-fibroblastic cells were identified based on their morphology, position in tissue and difference in GFP expression level. Namely, the *Acta2* model induced GFP expression in basal epithelial cells, vascular smooth muscle cells and pericytes; *Notch1* model induced GFP expression in subset of luminal epithelial cells and in endothelium.

### Mammary fragment transplantation

For transplantation experiments, mammary gland #4 from 5-week old *R26-mT/mG* females was harvested, spread on a 10 cm plastic petri dish and a 0.5×0.5×0.5 mm^3^ piece containing epithelial duct was dissected under a stereoscope using tdTomato fluorescence for guidance (Leica FM165C) and kept in CO2 independent medium on ice until use, but for maximum of 6 h. 3 weeks old nude females were anesthetized with isoflurane, after disinfection of abdominal area, reversed Y shaped incision was made into skin and #4 mammary glands were mobilized from the peritoneum. After cauterization of blood vessel in the connection of #5 and #4 mammary glands, proximal part of #4 mammary gland (from nipple to lymph node) was cut out. Small incision using iris scissors was made in the middle of the remaining fat pad to create a capsule, where a single piece of *R26-mT/mG* tissue was grafted. The skin opening was closed using bio-degradable suture and mice were carefully monitored until full recovery. During first 2 weeks after surgery, mice were supplied with analgesia (0.2 mg/ml of Ibuprofen in drinking water). 3.5 weeks after the surgery, the mice were euthanized and the mammary outgrowth was analyzed.

### Mammary organoid-fibroblast co-cultures

3D culture of mammary organoids and fibroblasts was performed as previously described^70^. Briefly, for isolation of primary mammary organoids (fragments), mammary glands were harvested, chopped and enzymatically digested [2 mg/ml collagenase A, 2 mg/ml trypsin, 5 μg/ml insulin, 50 μg/ml gentamicin (all Merck), 5% FBS (Hyclone/GE Healthcare) in DMEM/F12 (Thermo Fisher Scientific)] for 30 min at 37°C and 120 rpm. Resulting tissue suspension was treated with DNase I (20 U/ml; Merck) and enriched for organoids by 5 rounds of differential centrifugation at 450 × g for 10 s. The collagen/Matrigel gel matrix [MEM 1×, collagen 1 from rat tail (Corning) 2.5 mg/ml, 30% growth factor reduced Matrigel (Corning)] was mixed and pre-incubated for 1 h on ice before mixing with cells. 100 organoids, 80,000 FACS-sorted fibroblasts and 25 µl of collagen/Matrigel gel matrix were mixed well and plated on small patches of Matrigel on 8-well coverslip-bottom dishes (IBIDI) and overlayed with basal organoid medium (1× ITS, 100 U/ml of penicillin, and 100 μg/ml of streptomycin in DMEM/F12).

### Co-culture immunofluorescence

For immunofluorescent analysis of 3D co-cultures, the co-cultures were fixed with 4% paraformaldehyde for 40 min at RT, washed, and stored in PBS. The co-cultures were permeabilized with 0.5% Triton X-100 in PBS, blocked with 1% BSA, 5% FBS and 0.1% Triton X-100 in PBS (3D staining buffer, 3SB) and incubated with primary antibodies (supplementary table 2) in 3SB over 1-3 nights at 4°C. Then the co-cultures were washed for 3 h with 0.05% Tween-20 in PBS and incubated with fluorophore-conjugated secondary antibodies (ThermoFisher Scientific) and DAPI (1 μg/ml; Merck) in 3SB overnight at 4°C in dark. Then the co-cultures were washed for 3 h with 0.05% Tween-20 in PBS, cleared with 60% glycerol overnight at RT in dark. The co-cultures were imaged using LSM 880 (Zeiss).

### Statistical analysis

Statistical analysis was carried out in R using *stat_compare_means* function or *geom_signif* function of ggplot2 package. For comparing percentage, the *chisq.test* was used. N denotes number of samples used for statistical analysis, n denotes number of independent biological replicates, n.s. means not significant, *p < 0.05, **p < 0.01, ***p < 0.001, ****p < 0.0001. The box plots show median, the box borders are Q2-Q3, the whiskers are minimum to maximum, big points are predicted outliers, dots show single samples.

## Author contribution

JS conceptualized the work, performed the experiments, analyzed data and wrote original manuscript, ZSK and SF acquired funding, conceptualized and supervised the work and wrote the manuscript. RJ performed CellChat analysis of scRNAseq data. MMF contributed to experimental work and edited the manuscript.

## Funding

Czech Science Foundation (GAČR GA23-04974S) and Ministry of Education, Youth and Sports (ERC CZ LL2323 FIBROFORCE) to ZSK, Barrande Fellowship (Ministry of Education, Youth and Sports); FRM FDM202106013570; Brno PhD Talent Scholarship funded by the Brno City Municipality to JS. This work was supported by the French National Research Agency (ANR) grants number ANR-21-CE13-0047 and ANR-22-CE13-0009, the Medical Research Foundation FRM “FRM Equipes” EQU201903007821, the FSER (Fondation Schlumberger pour l’éducation et la recherche) FSER20200211117, the Association for Research against Cancer (ARC) label ARCPGA2021120004232_4874, the Worldwide Cancer Research Foundation # 24-0216 and by Labex DEEP ANR-Number 11-LBX-0044 to SF.

## Acknowledgement

The authors would like to acknowledge the Cell and Tissue Imaging Platform – PICT-IBiSA (member of France–Bioimaging – ANR-10-INBS-04) of the U934/UMR3215 of Institut Curie for help with light microscopy. We acknowledge the core facility CELLIM of CEITEC, supported by the Czech-BioImaging large RI project (LM2023050 funded by MEYS CR), for their support with obtaining scientific data presented in this paper. We acknowledge the Cytometry platform of Institut Curie, The platform of *in vivo* experimentation of Institut Curie and the animal unit of Masaryk University. We thank to Danijela M. Vignijevic and Pavel Tomancak for constructive discussions.

## Conflict of interest statement

The authors declare no conflict of interest.

## Notes

### Competing Interest Statement

The authors have declared no competing interest.

