## supplementary materials for "Contractile fibroblasts are recruited to the growing mammary epithelium to support branching morphogenesis"

### Supplementary figures

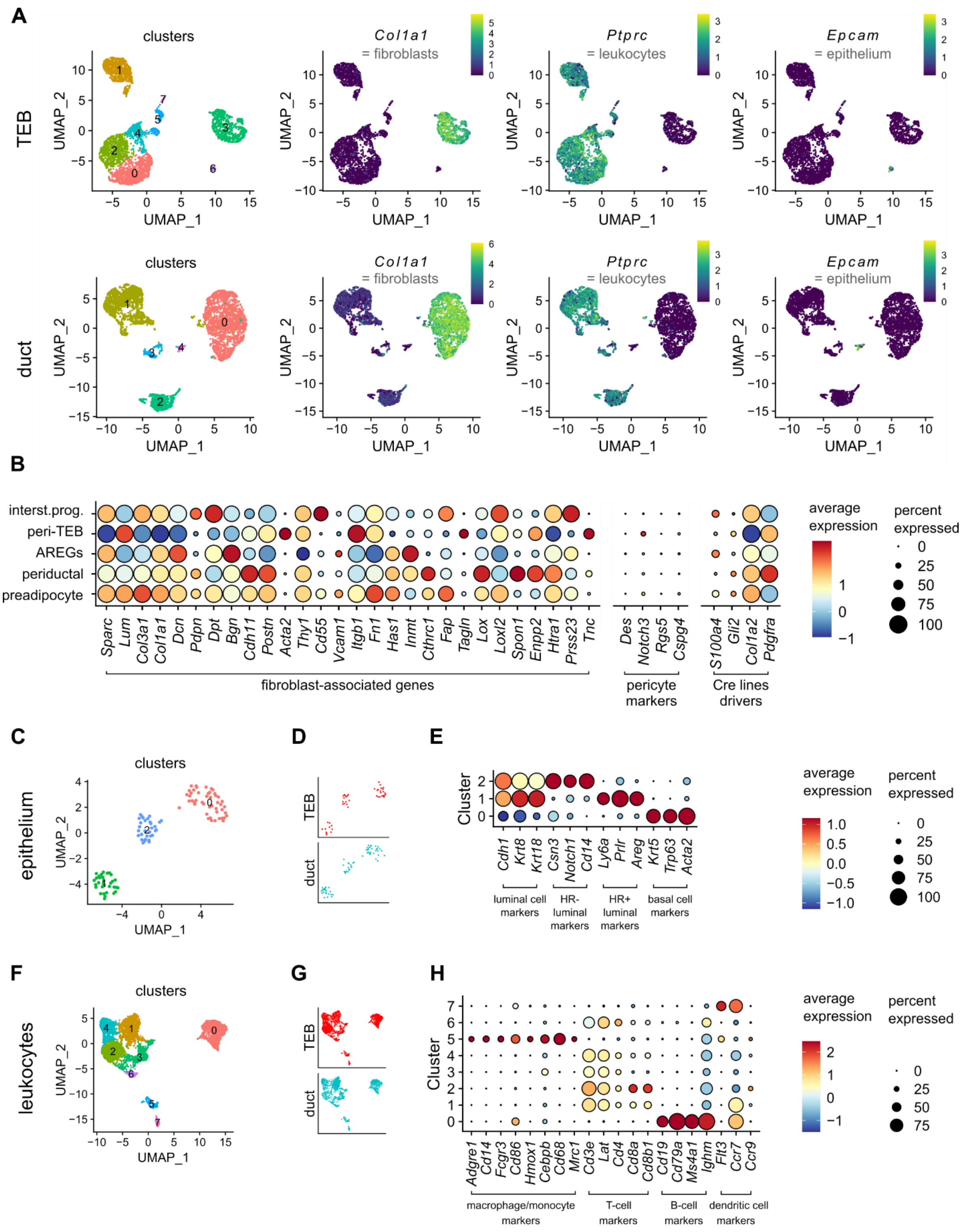

#### *Supplementary figure 1. scRNAseq of pubertal mammary gland*

**A.** UMAP plots showing cell clusters and canonical fibroblast, leukocyte and epithelial cell genes in the micro-dissected samples containing TEBs or ducts. **B.** Dot plot showing expression of fibroblast-associated genes, pericyte maker genes and genes previously used to drive Cre expression in mammary fibroblasts. **C-E.** Analysis of subsets of epithelial cells merged from the TEB and duct samples: UMAPs showing clusters (**C**) and cell origin (**D**), and a dot plot showing expression of epithelial cell type markers (**E**). **F-H.** Analysis of subset of leukocytes merged from TEB and duct sample: UMAPs showing clusters (**F**) and cell origin (**G**), and a dot plot showing expression of immune cell type markers (**H**).

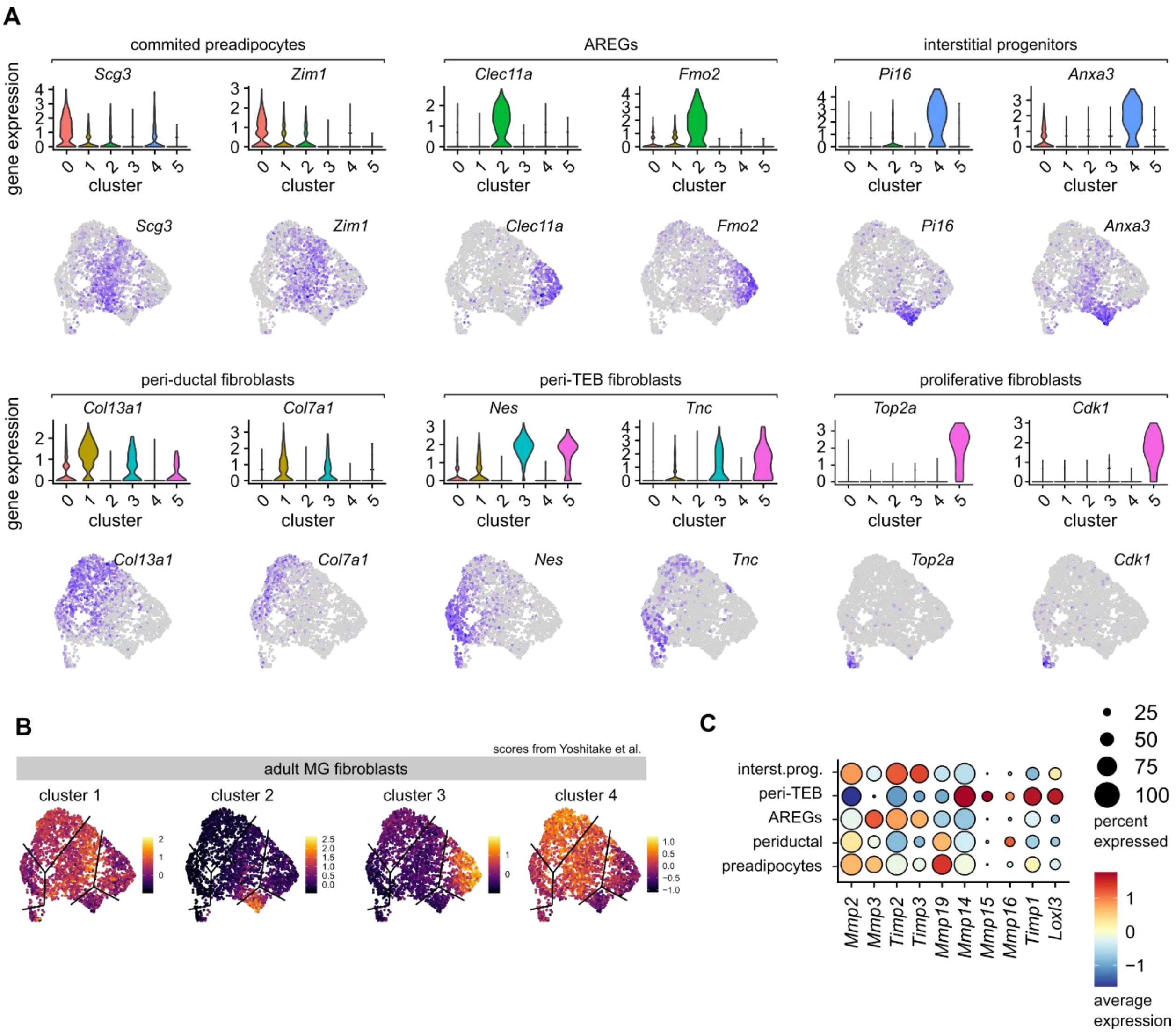


#### *Supplementary figure 2. scRNAseq analysis of pubertal mammary fibroblasts*

**A.** Violin plots and UMAPs showing expression of additional cluster marker genes. **B.** Transcriptomic scores of the 4 clusters of fibroblasts from adult mammary gland (MG) (Yoshitake et al. 2022) plotted over our dataset of mammary fibroblasts. The black lines indicate approximate borders between our 6 fibroblast clusters. **C.** Dot plot showing expression of ECM remodeling genes in our fibroblast clusters.

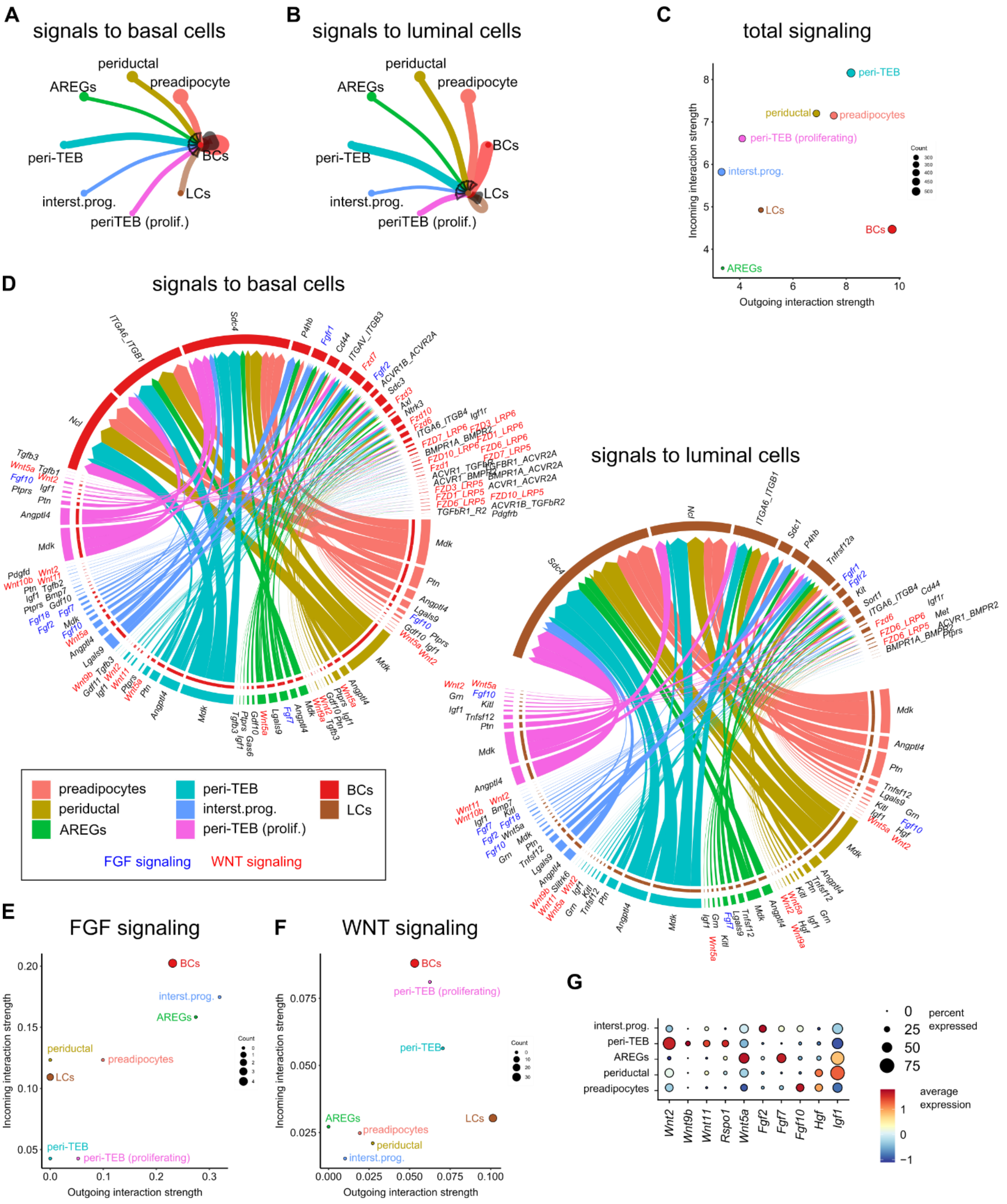


#### *Supplementary figure 3. CellChat analysis of pubertal mammary fibroblast-epithelium interactome*

**A, B.** Quantification of predicted signals originating in fibroblasts and directing into basal (BCs) (**A**) or luminal (LCs) (**B**) epithelial cells. The thickness of the line represents the signal strength.

**C.** A graph of overall incoming and outgoing signals for single fibroblast clusters and epithelial cells. **D.** Chord diagrams showing predicted signals incoming into basal or luminal epithelial cells from the identified fibroblast clusters. Members of FGF signaling are colored in blue, members of WNT signaling in red. **E, F.** Graphs showing the strength of incoming and outgoing signal of FGF (**E**) and WNT (**F**) signaling. **G.** A dot plot showing expression of selected signaling molecule genes in fibroblast clusters.

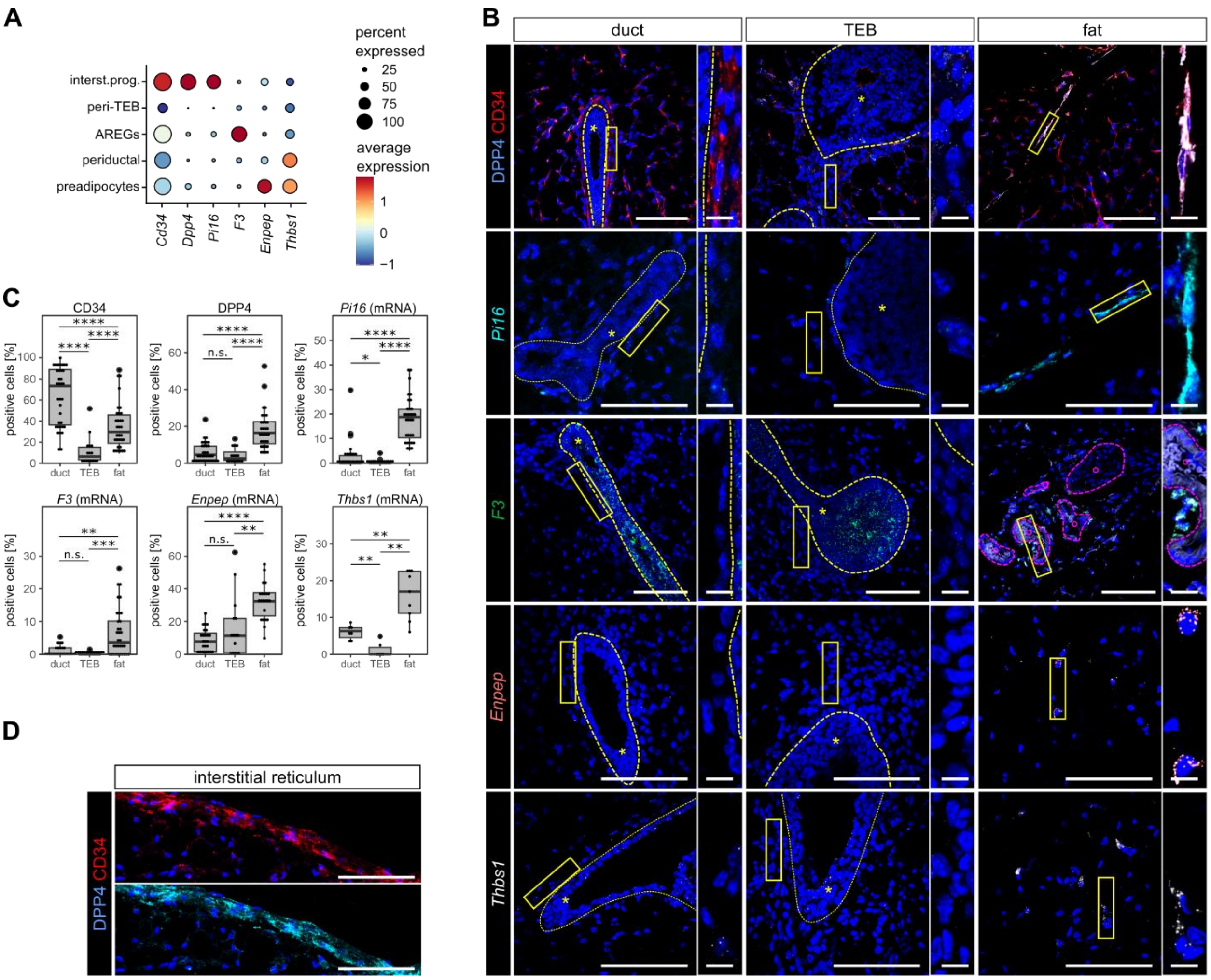


#### *Supplementary figure 4. Detection of fat pad clusters*

**A.** A dot plot showing expression of marker genes used for spatial mapping. **B.** Representative images showing expression of various gene or protein markers in regions containing duct, TEB or distal fat pad. The insets show details on fibroblasts. The dashed yellow line demarcates epithelial compartment, which is indicated by *. The dashed purple line demarcates blood vessel, which is indicated by °. Scale bar: 100 μm, 10 μm in detail. **C.** Quantification of marker-positive stromal cells in different regions of the mammary gland, shown as box plot. The dots show single fields of view (FOV), n = 3 biological replicates, N = 89 FOVs for CD34 and DPP4, 90 FOVs for *Pi16*, 74 FOVs for *F3*, 81 FOVs for *Enpep*, 28 FOVs for *Thbs1*, statistical analysis: Wilcoxon test. **D.** Representative images showing expression of DPP4 and CD34 in interstitial reticulum at the border of the mammary gland (D). Scale bar: 100 μm.

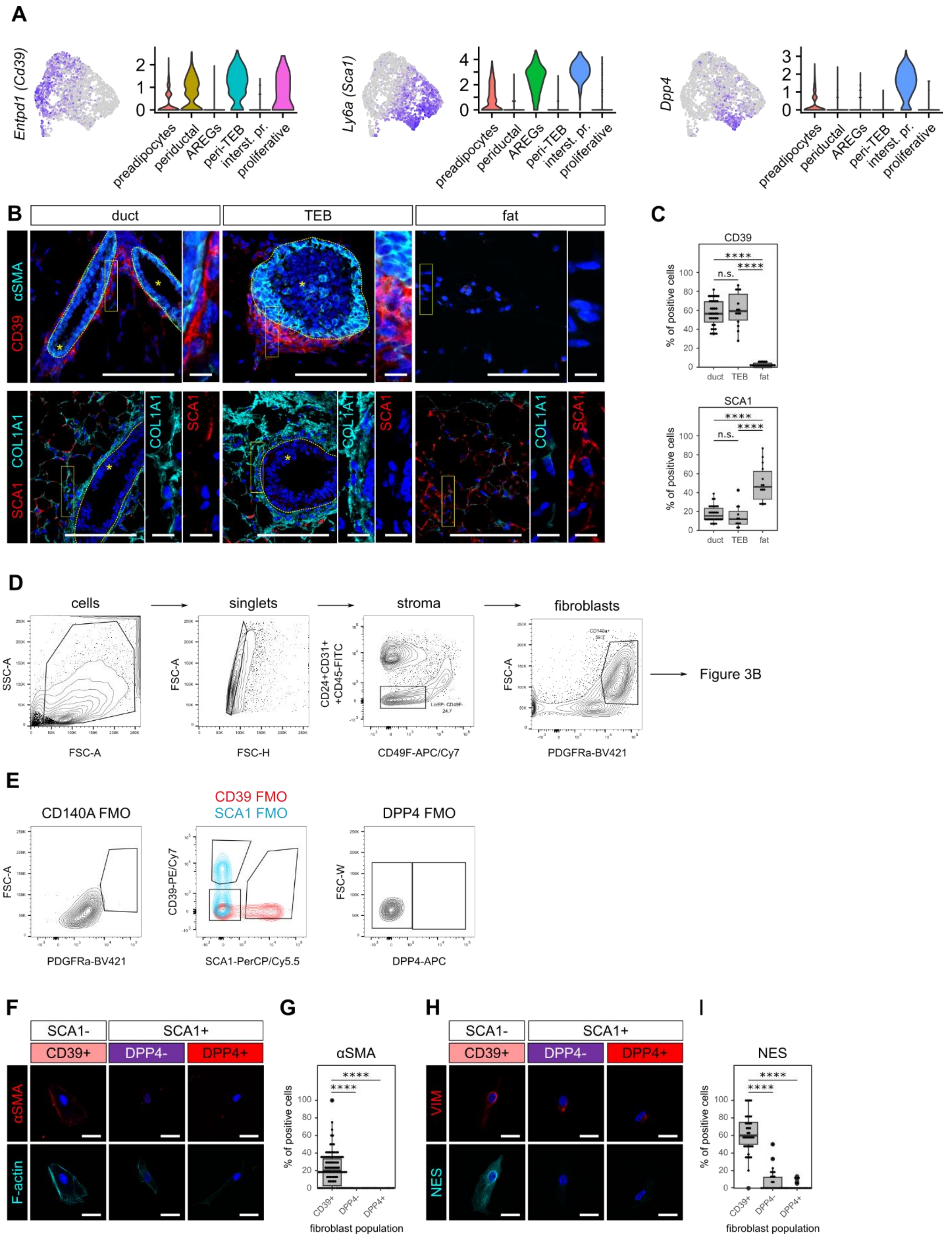
A
Entpd1 (Cd39)
Ly6a (Sca1)
Dpp4
preadipocytes
periductal
AREGs
peri-TEB
interst. pr.
proliferative
B
duct
TEB
fat
αSMA
CD39
COL1A1
SCA1
C
CD39
SCA1
% of positive cells
n.s.
****
duct
TEB
fat
D
cells
singlets
stroma
fibroblasts
SSC-A
FSC-A
FSC-H
CD24+CD31+ +CD45-FITC
CD49F-APC/Cy7
PDGFRa-BV421
Figure 3B
E
CD140A FMO
CD39 FMO
SCA1 FMO
DPP4 FMO
CD39-PE/Cy7
SCA1-PerCP/Cy5.5
FSC-W
DPP4-APC
F
SCA1-
SCA1+
CD39+
DPP4-
DPP4+
F-actin
G
H
VIM
NES
I
fibroblast population

#### *Supplementary figure 5. Flow cytometry analysis of mammary fibroblasts.*

**A.** UMAPs and violin plots showing expression of *Entpd1*, *Ly6a* and *Dpp4* in mammary fibroblasts. **B.** Representative images showing expression of CD39 and SCA1 in regions containing duct, TEB or distal fat pad. The insets show detail on fibroblasts. The dashed yellow line demarcates epithelial compartment, which is indicated by *. Scale bars: 100 µm, 10 µm in detail. **C.** Quantification of marker positive stromal cells in different regions of mammary gland, shown as box plot, the dots show single fields of view (FOV), n = 3 biological replicates, N = 68 FOVs for CD39 and 55 FOVs for SCA1, statistical analysis: Wilcoxon test. **D.** Representative flow cytometry scatter plots showing initial gating of isolated mammary gland cells, further gating of fibroblasts is shown in Figure 3A. **E.** Fluorescence minus one (FMO) controls for the experiment in D and Figure 3A. **F-I.** Characterization of peri-TEB marker expression. Representative images (**F**) and quantification (**G**) of αSMA expression in FACS-sorted populations of mammary fibroblasts. **H, I.** Representative images (**H**) and quantification (**I**) of NES expression in FACS-sorted populations of mammary fibroblasts. The dots of box plots in G and I represent single FOVs, n = 2 and 3 independent experiments for αSMA and NES, respectively; N = 848 cells in CD39+, 147 cells in DPP4- and 237 cells in DPP4+ population for αSMA; N = 479 cells in CD39+, 251 cells in DPP4- and 226 cells in DPP4+ population for NES. Scale bars: 10 µm.

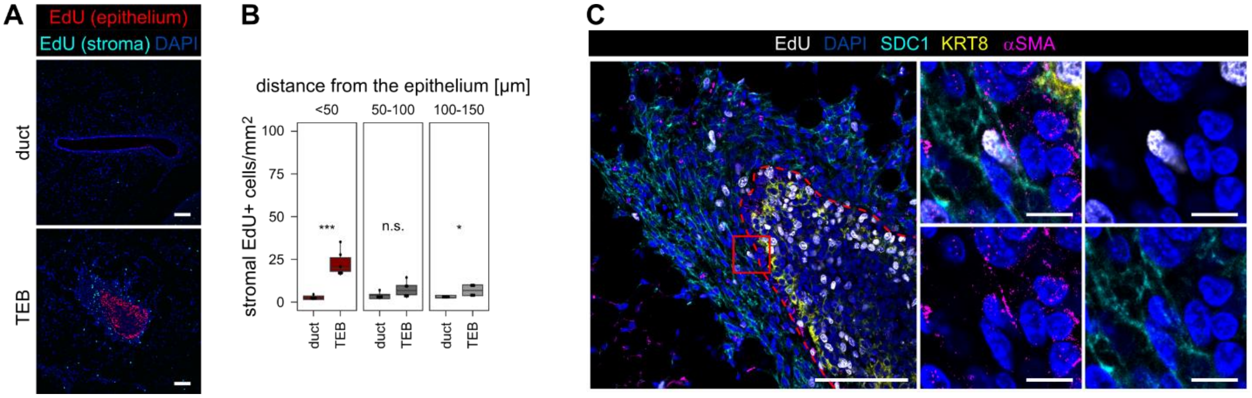


***Supplementary figure 6. Detection of proliferation in pubertal mammary stroma***

**A-C.** Detection of EdU incorporation after 2 h pulse on histological sections of the mammary gland. **A.** EdU signal is separated by manually drawing a mask in stromal and epithelial areas. Scale bar: 100 µm. **B.** Quantification of EdU+ stromal cells in relation to distance from the epithelial structures; n = 11 fields of view, statistical analysis: Wilcoxon test. **C.** Micrograph of the TEB from **A** showing an EdU+ αSMA+ SDC1+ peri-TEB fibroblast. Scale bar: 100 µm, 10 µm in detail.

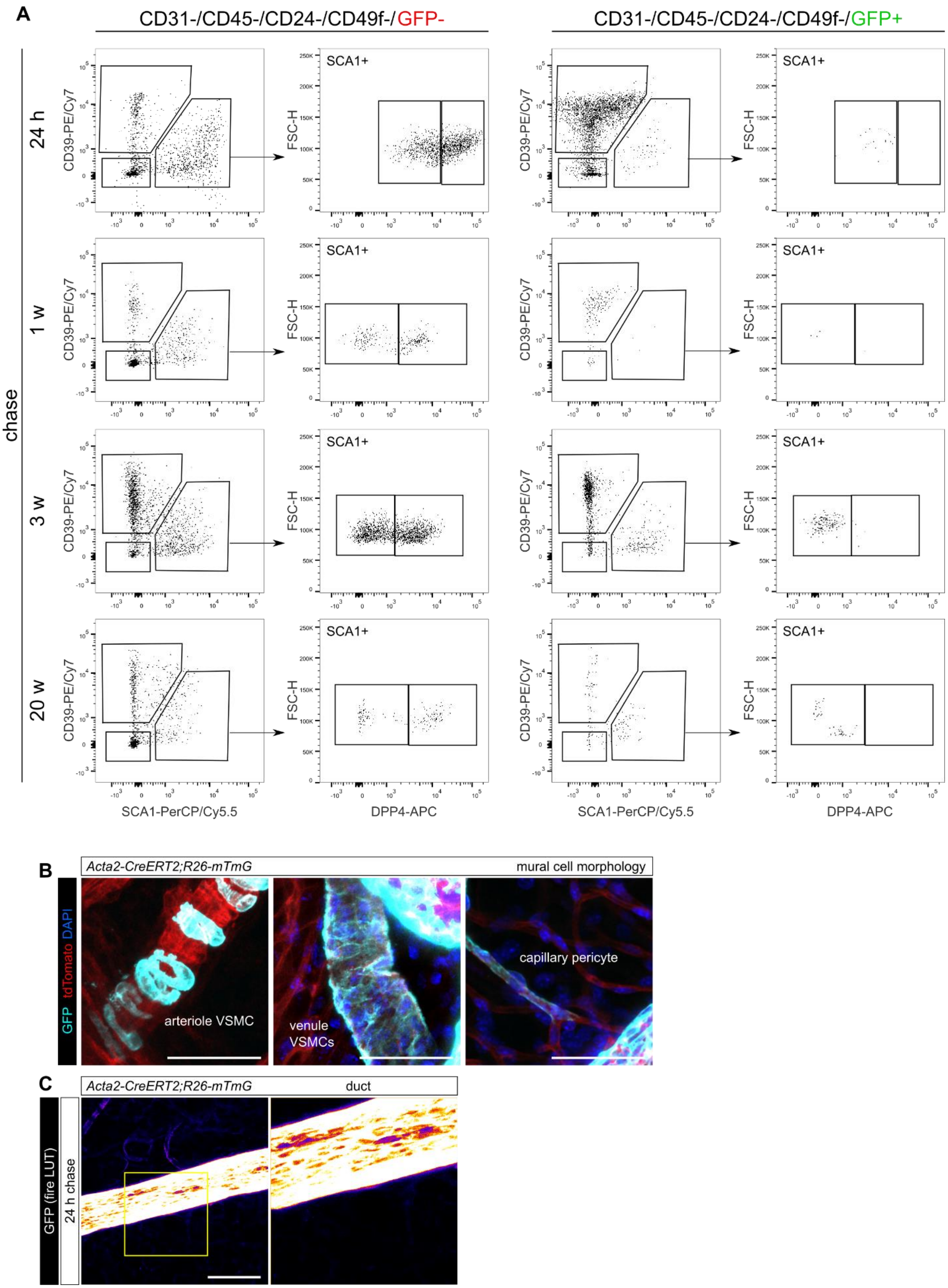
A
CD31-/CD45-/CD24-/CD49f-/GFP-
CD31-/CD45-/CD24-/CD49f-/GFP+
chase
24 h
1 w
3 w
20 w
SCA1+
CD39-PE/Cy7
FSC-H
SCA1-PerCP/Cy5.5
DPP4-APC
B
Acta2-CreERT2;R26-mTmG
mural cell morphology
GFP tdTomato DAPI
arteriole VSMC
venule VSMCs
capillary pericyte
C
Acta2-CreERT2;R26-mTmG
duct
GFP (fire LUT)
24 h chase

#### *Supplementary figure 7. Fibroblast tracing in the Acta2-CreERT2;R26-mTmG mouse model*

**A.** Representative flow cytometry scatter plots showing gating of fibroblast populations in GFP- and GFP+ stromal cells, used in quantification in Figure 3G. **B.** Representative images of mural cells visualized with the *Acta2-CreERT2;R26-mTmG*: vascular smooth muscle cells (VSMCs) and pericytes show characteristic shape and association with blood vessels. **C.** Z-stack and magnification of a duct from *Acta2-CreERT2;R26-mTmG* mouse induced with a 24 h pulse of tamoxifen (shown in Figure 3I). GFP channel is shown as “fire” lookup table (LUT). Scale bar: 100 µm.

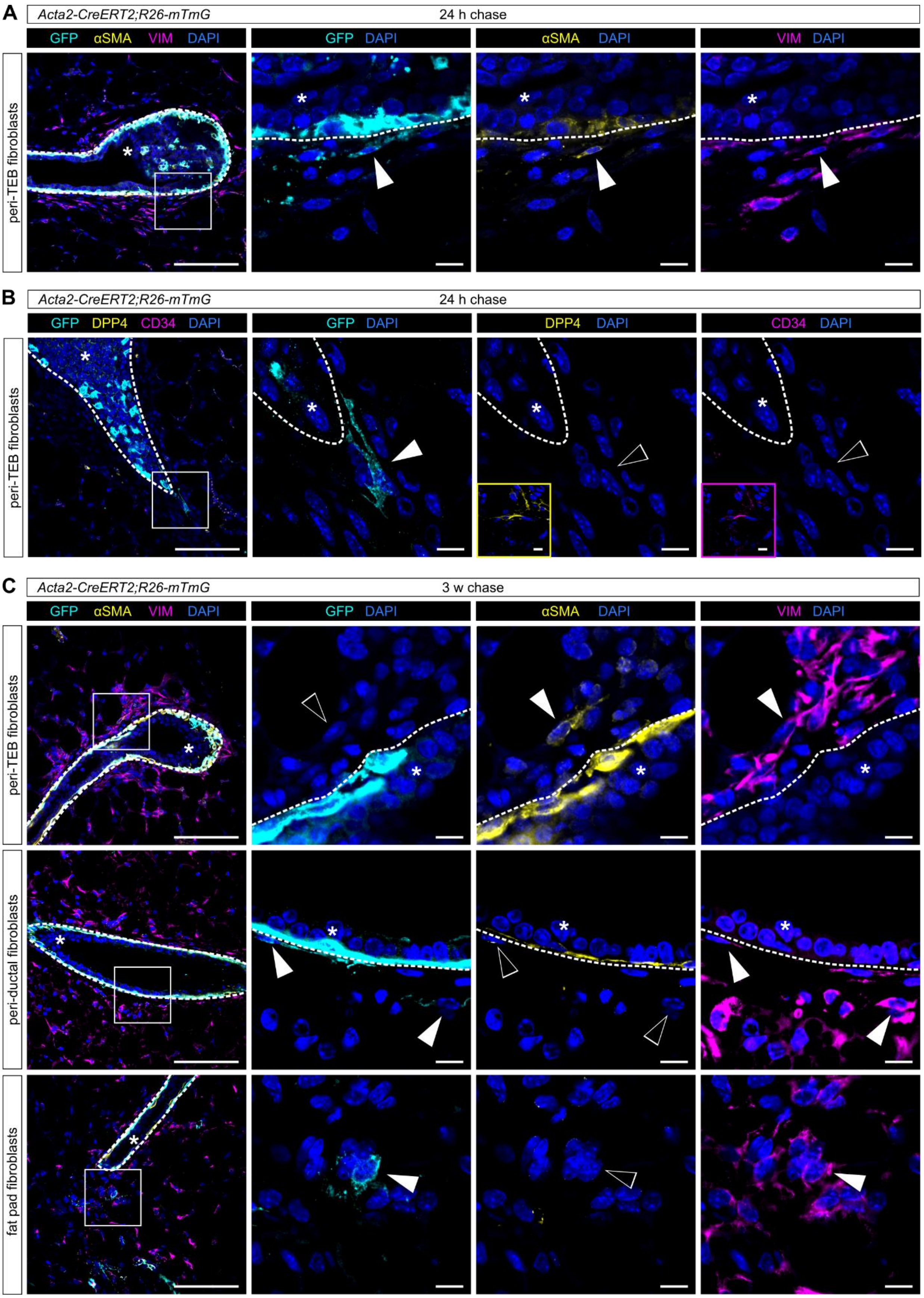
A
Acta2-CreERT2;R26-mTmG
24 h chase
GFP αSMA VIM DAPI
GFP DAPI
αSMA DAPI
VIM DAPI
peri-TEB fibroblasts
B
Acta2-CreERT2;R26-mTmG
24 h chase
GFP DPP4 CD34 DAPI
GFP DAPI
DPP4 DAPI
CD34 DAPI
peri-TEB fibroblasts
C
Acta2-CreERT2;R26-mTmG
3 w chase
GFP αSMA VIM DAPI
GFP DAPI
αSMA DAPI
VIM DAPI
peri-TEB fibroblasts
peri-ductal fibroblasts
fat pad fibroblasts

##### *Supplementary figure 8. Immuno-histological analysis of traced fibroblasts in the Acta2-CreERT2;R26-mTmG mouse model*

**A-C.** Representative images of mammary TEBs of an *Acta2-CreERT2;R26-mTmG* mouse induced with tamoxifen at 5 weeks of age and after a 24 h (**A, B**) or 3 week (**C**) chase, with details zooming on GFP+ fibroblasts, stained for VIM and αSMA (**A, C**) or CD34 and DPP4 (**B**). The yellow and magenta insets in **B** show positive staining of fat pad fibroblasts photographed in the same section. The dashed white line demarcates epithelial compartment, which is indicated by *. Full white arrowheads point out positive staining, empty arrowheads point out negative staining. Scale bars: 100 µm, 10 µm in detail.

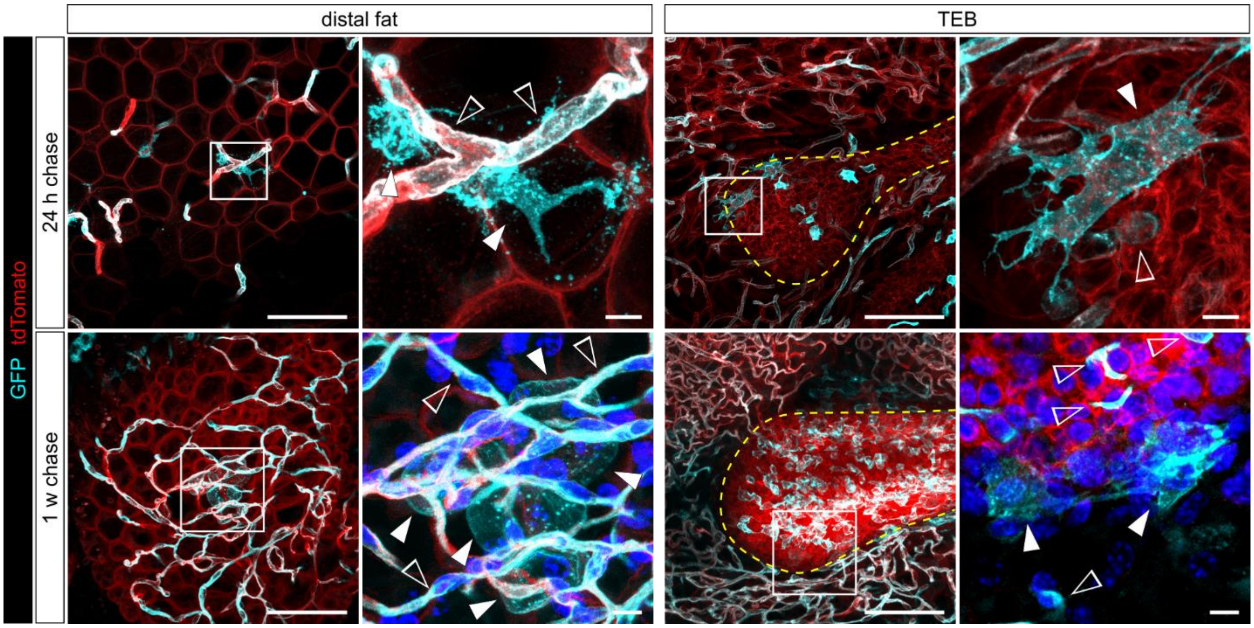


***Supplementary figure 9. Fibroblast visualization in the Notch1-CreERT2;R26-mTmG cleared mammary glands***

A distal fat pad region and a TEB region of a *Notch1-CreERT2;R26-mTmG* mice induced with a tamoxifen pulse and chased for 24 h (top panel) or 1 week (bottom panel). Full white arrowheads show GFP+ fibroblasts or adipocytes, empty arrowheads show GFP+ epithelial or endothelial cells. Scale bars: 100 µm, 10 µm in detail.

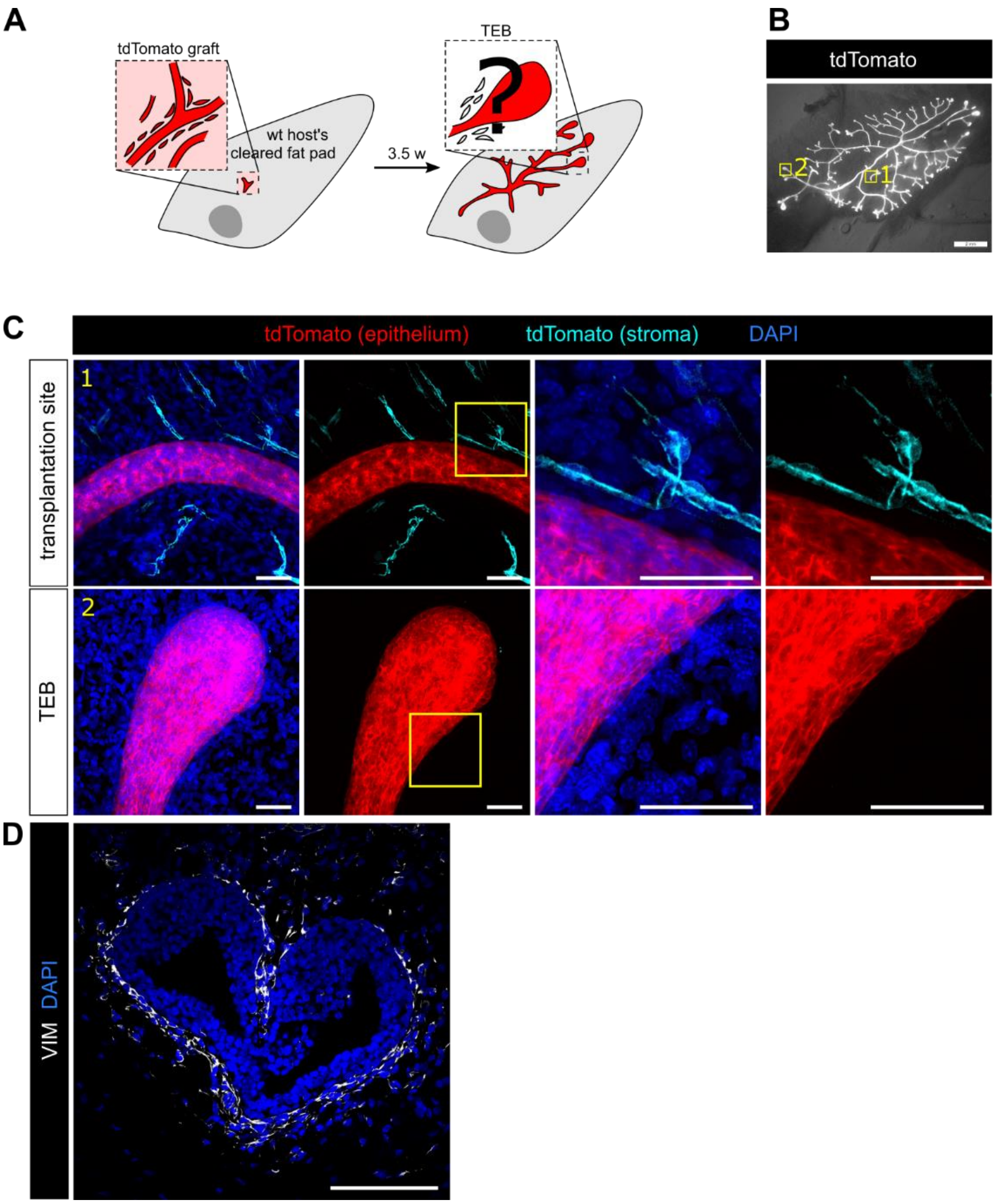


***Supplementary figure 10. tdTomato+ mammary fragment transplantation in wild-type host***

**A.** A scheme depicting the transplantation of a tdTomato+ tissue fragment containing epithelial duct with surrounding stroma in a wild-type (wt) host cleared fat pad. **B.** A representative outgrowth of a regenerated gland 3.5 w after transplantation. Areas marked by yellow squares are zoomed-in in **C**. Scale bar: 1 mm. **C**. Z-stacks of the transplantation site (1) and a TEB (2). The tdTomato signal was divided into epithelial (red) and stromal (turquoise) using manually drawn masks based on morphology. The yellow squares indicate magnified areas. Scale bars: 50 µm. **D.** Staining of VIM on histological section of a TEB of a transplanted mammary gland shows the abundant presence of fibroblasts around the TEB. Scale bars: 100 µm.

### Supplementary tables

**Supplementary table 1: the gene scores**

| Study | Cluster | Genes | | | | | | | | | |
|---|---|---|---|---|---|---|---|---|---|---|---|
| **Merrick et al. 2019** | **Group 1** | Ark1c18 | Cd55 | anxa3 | Pi16 | Sema3c | Gap43 | Bmp7 | Dect2 | Dpp4 | Wnt2 |
| **Merrick et al. 2019** | **Group 2** | Ggt5 | Fabp4 | Pparg | Vcam1 | Bmper | Cd36 | Rarres2 | Apoe | Icam1 | |
| **Merrick et al. 2019** | **Group 3** | Fmo2 | Vit | Ifitm1 | Gdf10 | Clec11a | F3 | | | | |
| **Yoshitake et al. 2022** | **Fib_0** | Pi16 | Anxa3 | Sema3c | Myoc | Igfbp5 | Akr1c18 | Cd55 | Smpd3 | Ackr3 | Dpp4 |
| **Yoshitake et al. 2022** | **Fib_1** | Fabp4 | Lpl | Col4a1 | Sparcl1 | Cxcl14 | Hmgcs1 | Col6a3 | Smoc2 | Col5a3 | Col4a2 |
| **Yoshitake et al. 2022** | **Fib_2** | Postn | Mfap4 | Scg3 | Penk | Tnc | Apoe | Enpp2 | Mdk | Srpx | Thbs1 |
| **Yoshitake et al. 2022** | **Fib_3** | Gdf10 | F3 | Gas6 | Cxcl12 | Cst3 | Tmem176b | Fmo2 | Inmt | Mgp | Il6 |

***Supplementary table 1. The genes used for transcriptomic scores in Figur 1 and Supplementary figure 2***

**Supplementary table 2: antibodies and probes used**

| target | manufacturer | Reference # | species | clone | concentration | conjugate |
|---|---|---|---|---|---|---|
| **whole-mount staining (CUBIC)** | | | | | | |
| **target** | **manufacturer** | **Reference #** | **species** | **clone** | **concentration** | **conjugate** |
| ACTA2 | Sigma | C6198 | mouse | 1A4 | 1/300 | Cy3 |
| **IHC-IF** | | | | | | |
| **target** | **manufacturer** | **Reference #** | **species** | **clone** | **concentration** | **conjugate** |
| COLA1A | Cell Signaling Technologies | #72026 | rabbit | E8F4L | 1/100 | N/A |
| VIM | Cell Signaling Technologies | #5741 | rabbit | D21H3 | 1/400 | N/A |
| ACTA2 | Sigma | A2547 | mouse | 1A4 | 1/300 | N/A |
| SDC1 | BD | #553712 | rat | 281-2 | 1/400 | N/A |
| NES | Chemicon | MAB353 | mouse | rat401 | 1/100 | N/A |
| MYH10 | BioLegend | #909901 | rabbit | polyclonal | 1/200 | N/A |
| TNC | R&D | MAB2138 | rat | #578 | 1/30 | N/A |
| DPP4 | Abcam | ab187048 | rabbit | EPR18215 | 1/200 | N/A |
| CD34 | Abcam | ab8185 | rat | MEC 14.7 | 1/50 | N/A |
| CD39 | Cell Signaling Technologies | #14481 | rabbit | E2X6B | 1/200 | N/A |
| KRT8 | DSHB | TROMA-I | rat | TROMA-I | 1/200 | N/A |
| GFP | Novus Biologicals | NB100-1614 | chicken | polyclonal | 1/300 | N/A |
| SCA1 | A Becton Dickinson co. | 553334 | rat | E13-161.7 | 1/100 | biotin |
| **ICC-IF** | | | | | | |
| **target** | **manufacturer** | **Reference #** | **species** | **clone** | **concentration** | **conjugate** |
| ACTA2 | Sigma | A2547 | mouse | 1A4 | 1/300 | N/A |
| VIM | Cell Signaling Technologies | #5741 | rabbit | D21H3 | 1/200 | N/A |
| NES | Chemicon | MAB353 | mouse | rat401 | 1/100 | N/A |
| phalloidin | Sigma | | N/A | N/A | 1/300 | AF488 |
| **IF organoids** | | | | | | |
| **target** | **manufacturer** | **Reference #** | **species** | **clone** | **concentration** | **conjugate** |
| ACTA2 | Sigma | C6198 | mouse | 1A4 | 1/300 | Cy3 |
| ACTA2 | Sigma | A2547 | mouse | 1A4 | 1/300 | N/A |
| **flow cytometry/FACS** | | | | | | |
| **target** | **manufacturer** | **Reference #** | **species** | **clone** | **concentration** | **conjugate** |
| CD45 | BD | #561037 | rat | 30-F11 | 1/200 | APC/Cy7 |
| CD31 | BioLegend | #102439 | rat | 390 | 1/200 | APC/Cy7 |
| CD24 | BioLegend | #101849 | rat | M1/69 | 1/200 | APC/Cy7 |
| CD49f | BioLegend | #313628 | rat | GoH3 | 1/200 | APC/Cy7 |
| CD140A | BioLegend | #135923 | rat | APA5 | 1/50 | BV421 |
| CD39 | eBioscience | 25-0391-80 | rat | 24DMS1 | 1/200 | PE/Cy7 |
| SCA1 | BioLegend | #108123 | rat | D7 | 1/200 | PerCP/Cy5.5 |
| DPP4 | BioLegend | #137807 | rat | H194-112 | 1/200 | APC |
| **ISH probes** | | | | | | |
| **Target** | **Manufacturer** | **Reference #** | **channel** | | | |
| *Mfap4* | ACD | 421391 | C1 | | | |
| *Tnc* | ACD | 465021-C2 | C2 | | | |
| *Pi16* | ACD | 451311-C3 | C3 | | | |
| *F3* | ACD | 448691 | C1 | | | |
| *Enpep* | ACD | 862211-C3 | C3 | | | |
| *Thbs1* | ACD | 457891-C3 | C3 | | | |

***Supplementary figure 2. Antibodies and probes used***
